# Functional Dichotomy of Developmental Foxp3^+^ Treg Cell Subsets in the Visceral Adipose Tissue of Lean and Obese Mice

**DOI:** 10.1101/2025.03.12.642664

**Authors:** Acelya Yilmazer, Anne Eugster, Dimitra Maria Zevla, Sara Salome Helbich, Marie Boernert, Basak Torun, Elda Marsela, Emre Kirgin, Andreas Dahl, Andreas Petzold, Olivia Kershaw, Vasileia Ismini Alexaki, Antonios Chatzigeorgiou, Michael Delacher, Susan Schlenner, Karsten Kretschmer

## Abstract

Chronic inflammation and loss of Foxp3^+^ regulatory T (Treg) cells in the visceral adipose tissue (VAT) are hallmarks of the pathogenesis of insulin resistance and obesity. This study explores the roles of VAT Treg cells from thymic (tTreg) and peripheral (pTreg) developmental origin, revealing their opposing roles in metabolic inflammation. Obesity destabilized VAT tTreg cells, causing them to clonally expand into obesogenic Foxp3^−^IFN-γ^+^ T effector cells, enhancing pro-inflammatory type 1 responses. Genetic tTreg ablation prevented this shift, promoting anti-inflammatory type 2 response, reduced body weight, and improved insulin resistance. Compared to their tTreg counterpart, pTreg cells were functionally well adapted to maintain VAT homeostasis and protect against obesity. Genetic pTreg ablation promoted spontaneous obesity symptoms even with physiological calorie intake, and worsened VAT inflammation and liver steatosis on a high-calorie diet. These findings highlight tTreg instability as a pathogenic threat and pTreg cells as crucial regulators of metabolic homeostasis.

**Highlights:**

- VAT Tregs of lean mice originate from both thymic and peripheral Treg development
- High-calorie diet destabilizes tTregs that clonally expand into obesogenic IFN-γ^+^ Th1 cells
- Genetic tTreg deficiency improves steady-state metabolism and prevents diet-induced obesity
- Genetic pTreg deficiency promotes obesity in both sexes even with normal calorie intake
- VAT pTregs are particularly adapted to regulate VAT homeostasis, including adipogenesis

**In Brief:** Obesity and type 2 diabetes are characterized by insulin resistance, regulatory T (Treg) cell loss, and chronic inflammation in visceral adipose tissue (VAT). In this context, Yilmazer *et al.* dissect the functional roles of tTreg and pTreg cells. They show that VAT pTreg cells are particularly adapted to exert non-redundant homeostatic functions, and that pTreg deficiency predisposes to obesity even with normal calorie intake. In contrast, VAT tTreg cells can contribute to local inflammation by dedifferentiating into Foxp3^−^ Th1-polarized effector cells.

**Graphical abstract:** 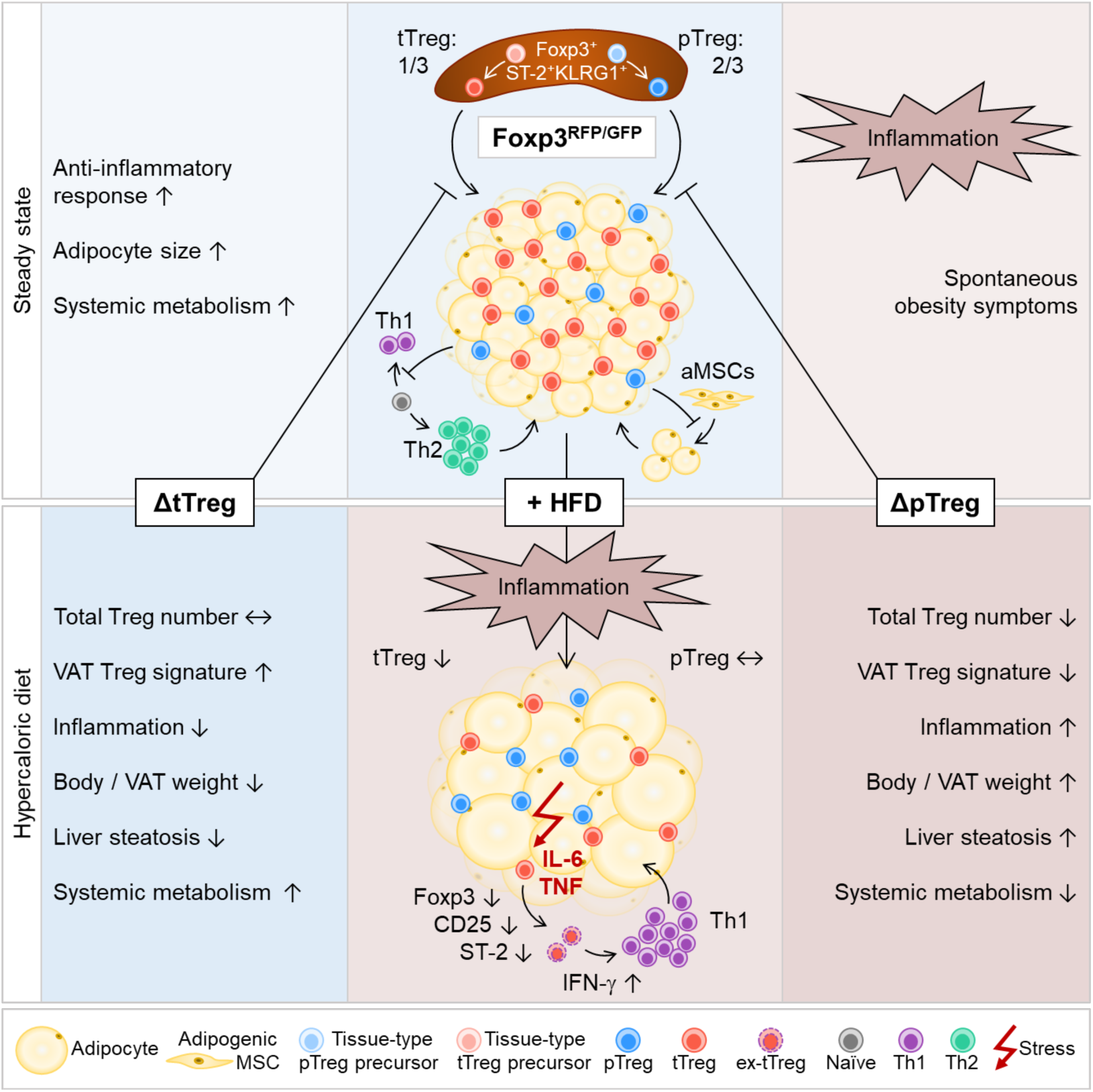

## INTRODUCTION

Obesity is a major global health challenge associated with chronic low-grade inflammation, particularly in visceral adipose tissue (VAT).^1–3^ This inflammation is a key driver of numerous metabolic complications, including insulin resistance, type 2 diabetes (T2D), cardiovascular diseases, and fatty liver disease.^4,5^ The importance of the metabolic and immunologic interplay in VAT for systemic metabolic homeostasis has been highlighted by surgical VAT removal that prevents obesity-induced multiorgan insulin resistance in mice.^6^ Among the immune cells that regulate VAT inflammation, CD4^+^ regulatory T (Treg) cells expressing the lineage specification factor Foxp3 play a central role.^7–9^ VAT Foxp3^+^ Treg cells exhibit a unique phenotype and functional properties compared to their counterparts in other tissues, including expression of the transcription factor PPAR-γ and the IL-33 receptor ST-2. In the lean state, VAT Treg cells are abundant and contribute to tissue homeostasis by suppressing pro-inflammatory responses.^7,10–12^ VAT Treg cells have also been implicated in non-immune functions by restraining differentiation of stromal adipocyte precursors via the oncostatin-M signalling pathway, thereby promoting insulin sensitivity and metabolic homeostasis.^13^ However, the manifestation of obesity correlates with a significant reduction in Treg abundance and function in VAT,^7,14–17^ which may have implications for pathogenesis, although the underlying mechanisms are poorly understood. The selective loss of Treg cells in VAT is also associated with an increase in pro-inflammatory immune cells, including M1 macrophages and T effector (Teff) cells, which exacerbates obesogenic VAT inflammation and metabolic dysfunction.^7,10,11^

Understanding the mechanisms that regulate VAT Treg cell abundance and function is crucial for developing therapeutic strategies aimed at mitigating obesity-associated inflammation and its metabolic sequelae. With regard to VAT Treg cell function, their developmental heterogeneity is a particularly important, albeit unresolved issue. The physiologic Treg cell compartment in lymphoid tissues consists of intrathymically (tTreg) and peripherally (pTreg) induced Treg cells that originate from distinct CD4^+^Foxp3^−^ precursor cells.^18,19^ Both tTreg and pTreg cell lineages are thought to synergistically act to maintain immune tolerance and prevent catastrophic autoimmunity.^20^ VAT Treg cells, like other ST-2^+^ tissue Treg cells, arise from lymphoid tissue-residing CD4^+^Foxp3^+^ common precursors and accumulate in non-lymphoid tissues, where they undergo further adaptations to fulfill homeostatic and regenerative functions.^21–23^ This includes the acquisition of PPAR-γ expression,^11^ which is crucial for the local accumulation and maintenance of Foxp3^+^ST-2^+^ Treg cells in VAT, as well as their specialized functions in regulating the immune and metabolic microenvironment. Unsurprisingly, the relative contribution of pTreg and tTreg cells to the VAT-Treg cell compartment has been the subject of investigations, although hampered by the lack of definite markers to faithfully distinguish between the two developmental subsets and the enormous phenotypic plasticity with which Treg cells are endowed.^24–26^ These studies therefore relied on experimental approaches, such as T cell receptor (TCR) repertoire analyses,^7,16^ the use of Nrp-1 and Helios as putative tTreg cell markers^27^ or the Foxp3^+^ Treg cell conversion of adoptively transferred conventional CD4^+^ T cells.^16,27,28^ The results revealed that the VAT Treg cell compartment is seeded from the thymus during the first weeks of life, followed by local proliferative expansion of particular tTreg cell clonotypes.^27,28^ In particular, studies on the TCR repertoire suggested that pTreg cells could also contribute to the VAT Treg cell compartment. However, the local recruitment of pre-formed pTreg cells or their *de novo* induction *in situ* has been difficult to demonstrate, which could potentially be attributed to the rather small proportion of pTreg cells in the total Treg cell pool.^25,29^ A functional role of pTreg cells in VAT has not been demonstrated yet,^7,27,28^ whereas the transcriptional profile of lymphoid tissue pTreg cells of dual Foxp3^RFP/GFP^ reporter mice shows striking similarities^25^ with that of total VAT Treg cells.^21,30^ This includes the upregulation of genes encoding ST-2, KLRG1, and IL-10, suggesting a possible relatedness between the two Treg cell populations.

These findings indicate that both pTreg and tTreg cells are involved in the regulation of VAT homeostasis, but leave open the question of their relative contribution, mechanisms of action and responses to obesity-induced stressors. To define the roles of pTreg and tTreg cells *in vivo* in loss-of-function studies, we previously generated complementary mouse lines based on dual Foxp3^RFP/GFP^ reporter mice^25^ selectively deficient in pTreg (ΔpTreg) or tTreg (ΔtTreg) cell development.^29,31^ These mouse genetic tools have been proven useful in demonstrating that tTreg cells are key regulators of pancreatic β-cell autoimmunity.^29^ Here, we used Foxp3^RFP/GFP^ mice and their ΔpTreg and ΔtTreg derivatives, in conjunction with multicolor flow cytometry and single-cell transcriptomics, to mechanistically decipher the function of pTreg and tTreg cells in VAT of lean and obese mice.

## RESULTS

### Obesity selectively reduces the thymus-derived Treg cell subset in visceral adipose tissue

The reduction in the number of total VAT Treg cells is a hallmark of obesity that has also been implicated in its pathogenesis.^7,10,11,15^ To dissect if and how pTreg and tTreg cells are affected, we first examined the effects of high fat diet (HFD) in kinetic studies in genetically non-modified wild-type (WT) and dual Foxp3^RFP/GFP^ reporter mice on the C57BL/6 background (Figure 1A). Compared to normocaloric diet (ND), feeding of HFD for up to 20 weeks resulted in significant increases in body weight (Figure 1B and 1C), adipose tissue (AT) weight (Figure 1D), and total cell counts of the non-adipocyte stromal vascular fraction (SVF) obtained from ATs (Figure 1E). The proportion of CD45^+^ hematopoietic lineage cells of the SVF were also significantly increased [VAT: ND, 35.8 ± 6.0%; HFD, 57.0 ± 13.5%; subcutaneous AT (SAT): ND, 31.9 ± 12.6%; HFD, 48.6 ± 8.0%]. Substantial changes in innate immune cells (Figures 1F and S1A) [*i.e.*, pro-inflammatory M1 macrophages (CD64^+^CD11b^+^F4/80^+^-gated CD11c^high^CD206^low^) and inflammatory dendritic cells (iDCs; CD64^⎻^CD11c^+^- gated CD11b^+^CD103^−^) ↑; anti-inflammatory M2 macrophages (CD64^+^CD11b^+^F4/80^+^-gated CD11c^low^CD206^high^) and eosinophils (EOs; CD64^⎻^CD11b^+^-gated Siglec-F^+^F4/80^+^): ↓] reflected the expected transition from an anti-inflammatory type 2 to a pro-inflammatory type 1 immune response in the VAT of HFD-fed mice.^32–35^ At 20 weeks of feeding, we found HFD to increase the population size of VAT CD8^+^ T cells, but we did not observe such changes in CD4^+^ T cells at any of the time points examined (Figures S1B, S2A, 1G). We observed, however, the expected HFD-induced reduction of the total Foxp3^+^ Treg cell compartment among CD4-gated cells (Figures 1G and 1H), which in Foxp3^RFP/GFP^ mice could be entirely attributed to tTreg cells co-expressing RFP and GFP (ND: 24.4 ± 4.0%; HFD: 8.7

**Figure 1.**
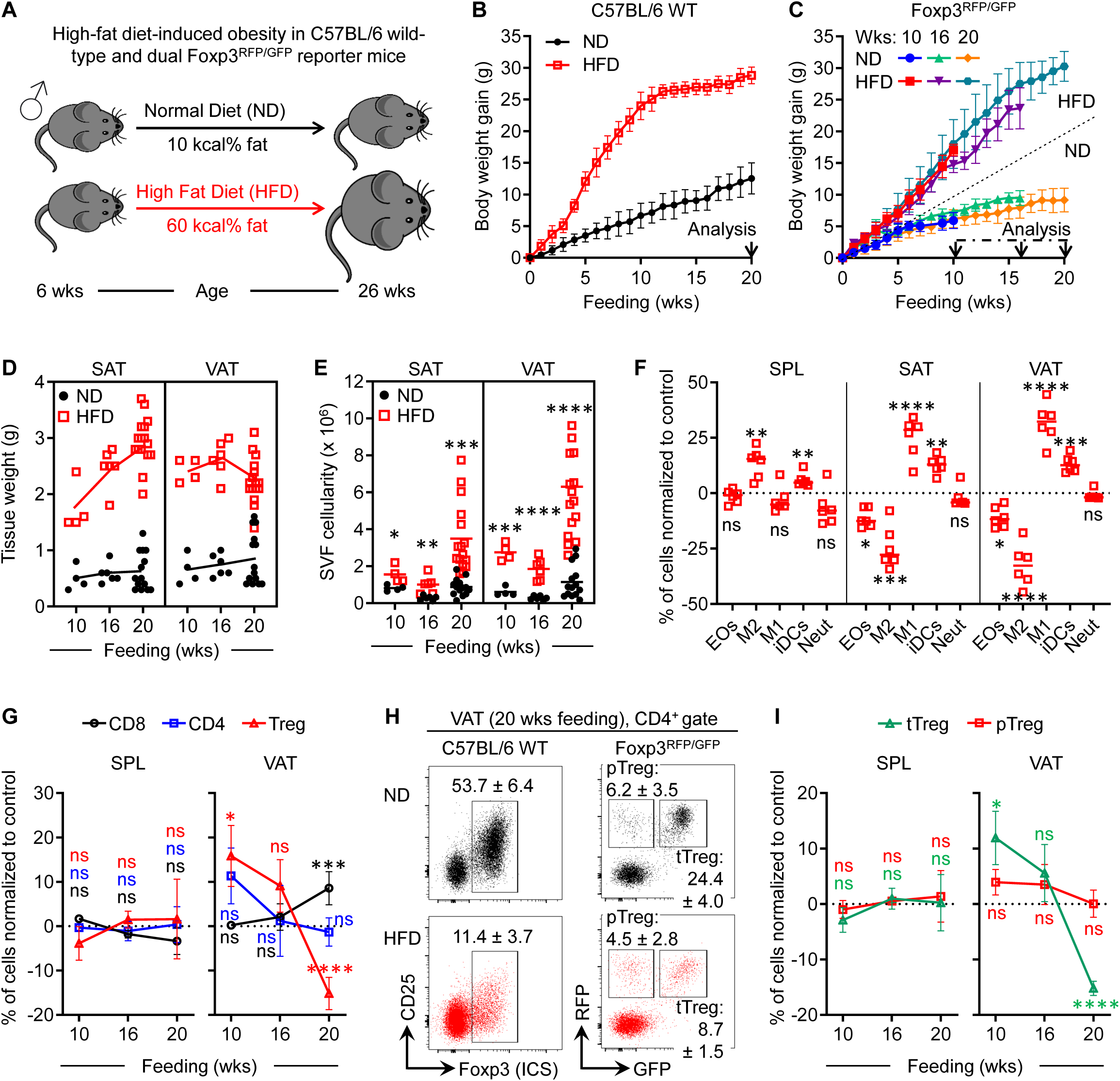
HFD selectively reduces thymus-derived VAT Treg cells (A) Experimental scheme of HFD-induced obesity. (B) Body weight gain in wild-type mice. (n = 6 – 8/group). (C-I) HFD-induced obesity in Foxp3^RFP/GFP^ mice. (C) Body weight gain. (n = 4 – 16/group). Arrowheads indicate analysis time points. (D) Weight of subcutaneous (left) and visceral (right) adipose tissue. (E) SVF cell count in subcutaneous (left) and visceral (right) adipose tissue. (F) Obesity-induced shift to proinflammatory type 1 immunity in VAT. Cell frequencies after 16 weeks of HFD feeding, normalized to mean frequencies in ND-fed mice. (G) Loss of VAT Treg cells. Frequencies of conventional CD8^+^ and CD4^+^ T cells, along with RFP^+^ Treg cells, in HFD-fed Foxp3^RFP/GFP^ mice, normalized to mean frequencies under ND feeding. (H) Representative flow cytometry of VAT Treg cells in WT (left) and Foxp3^RFP/GFP^ (right) mice on ND (top) or HFD (bottom). (I) Selective loss of tTreg cells in VAT. Frequencies of tTreg and pTreg cells in HFD-fed Foxp3^RFP/GFP^ mice normalized to mean frequencies in ND-fed mice. Mean ± SD (B, C, G, H, I). Symbols in (D, E, F) represent individual mice. Data represent one of three independent experiments (n = 3 - 6 mice). Unpaired t-test: ns, not significant; *p ≤ 0.05, **p < 0.01, ***p < 0.001, ****p < 0.0001 (E, F, G, I). See also Figures S1 and S2.

± 1.5%) (Figure 1H, right panels). This selective tTreg cell loss in VAT occurred late in the HFD feeding period and was not observed in the SAT or spleen (Figures 1I and S2B). This resulted in a significant shift in the Treg-to-immune effector cell ratio in VAT, disadvantaging tTreg cells at 20 weeks of HFD feeding (Figure S2C). Normalizing cell counts to tissue weight (Figure S2D) revealed a numerical increase across all examined immune cell compartments within the first 10 weeks of HFD feeding. However, by 20 weeks, the total CD45^+^ immune cell population, including CD4^+^ T cells and pTreg cells (RFP^+^GFP^−^), exhibited a marked expansion, whereas tTreg cells showed no comparable increase (Figure S2D). Interestingly, we did not observe significant differences in the size of the pTreg cell compartment in either ND- or HFD-fed mice at any of the time points examined (Figures 1H, 1I, and S2B), indicating that VAT pTreg cells failed to colonize the niche after selective tTreg cell loss. In mice with constitutive tTreg cell deficiency, however, the pTreg cell compartment significantly increased in size (see below). To explore a causal relationship between the HFD-mediated manifestation of obesity symptoms and the concomitant loss of tTreg cells in the VAT, we next conducted loss-of-function studies.

### Selective tTreg cell paucity ameliorates HFD-induced obesity

To examine the functional relevance of selective tTreg cell loss observed in the obese VAT (Figures 1H, 1I, and S2B), we utilized Foxp3^RFP/GFP^ mice deficient in tTreg cells (ΔtTreg mice) due to their ablation during thymic development, while sparing the development of pTreg cells.^29,31^ In C57BL/6 ΔtTreg mice, the pTreg cell sister population functionally compensates for the lack of tTreg cells and prevents the manifestation of autoimmune diseases^29^ commonly observed in models with complete deficiency of all Foxp3^+^ Treg cells.^36,37^ Compared to littermate controls, selective tTreg cell deficiency (Figure 2A) significantly attenuated HFD-induced body weight gain (Figure 2B). In these experiments, the body weight of both ΔtTreg and Foxp3^RFP/GFP^ mice was comparable at the beginning of HFD feeding at 6 weeks of age (Figure 2C). Foxp3^RFP/GFP^ mice showed a continuous body weight gain until the end of the feeding period with the expected kinetics (see Figures 1B and 1C for comparison). However, the body weight gain of ΔtTreg mice increasingly slowed down from the third week of feeding (Figure 2B). Approximately ⅓ of ΔtTreg mice (Figure 2C) failed to show any HFD-induced body weight increase even after ≥ 15 weeks of feeding, compared to age-matched Foxp3^RFP/GFP^ and ΔtTreg mice fed a normocaloric vegetarian standard diet (VegND). While many HFD-fed ΔtTreg mice additionally showed attenuated increases in AT weight (Figure S3A) and total SVF cellularity (Figure S3B), these data suggest that the impact of selective tTreg cell paucity on HFD-induced body weight gain extends beyond VAT and SAT, and may include other fat tissues. The proportional distribution of innate immune cells (Figures S3C and S3D) indicated that transition from anti-inflammatory type 2 to pro-inflammatory type 1 immunity in the SAT and VAT was reverted in HFD-fed ΔtTreg mice.

**Figure 2.**
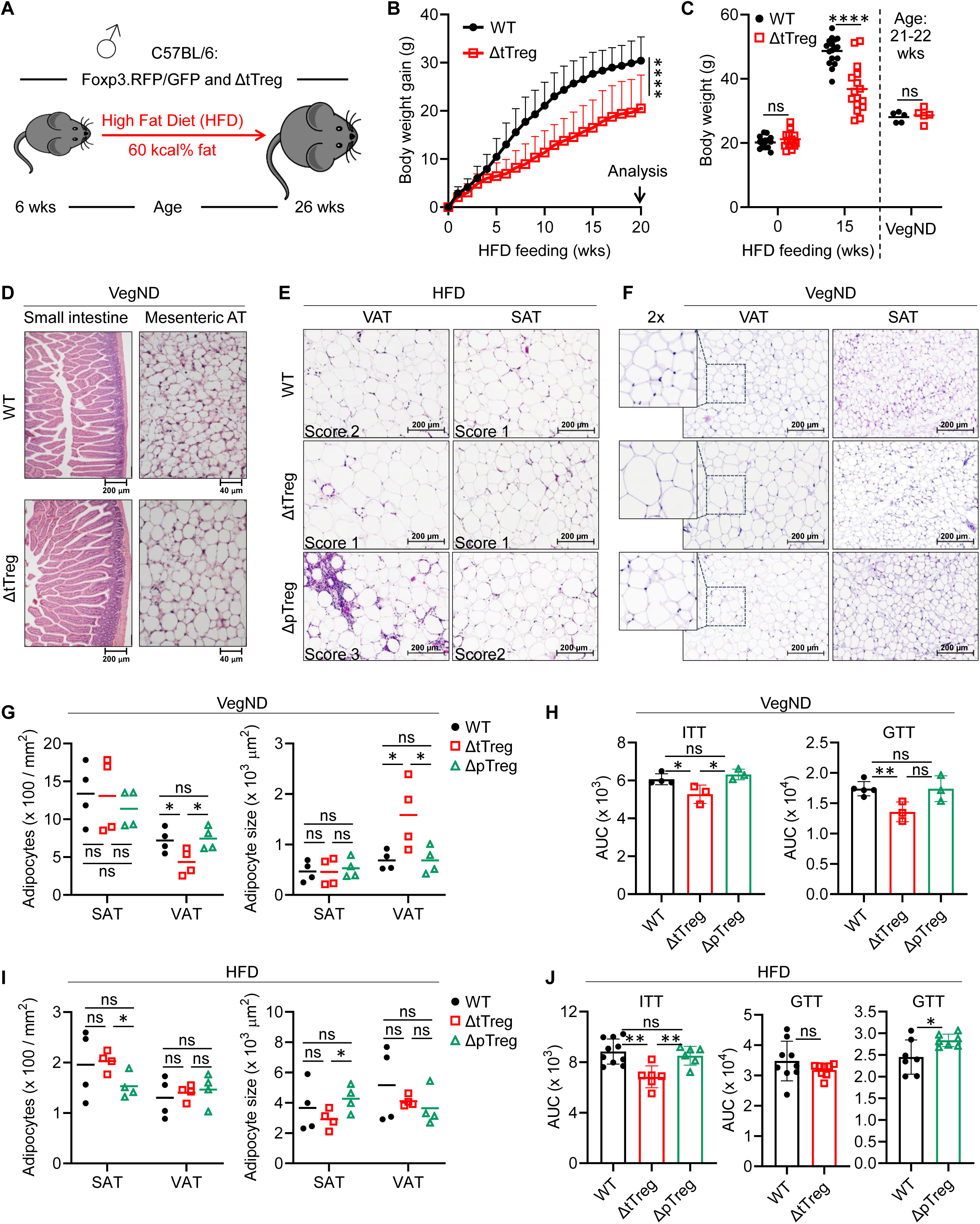
Genetic tTreg cell ablation improves clinical signs of obesity and metabolic fitness (A-C) Attenuated HFD-induced body weight gain in ΔtTreg mice. (A) Experimental scheme of HFD feeding. (B) Body weight gain of Foxp3^RFP/GFP^ (WT, n = 15) and ΔtTreg (n = 13) mice. Mean ± SD. Two-way ANOVA with Sidak’s correction; p-values in Figure 1. See also Figure S3. (C) Body weight at baseline (0 weeks) and 15 weeks of HFD feeding (WT, n = 17; ΔtTreg, n = 16), and of age-matched, VegND-fed mice (n = 5/group). Unpaired t-test: p-values in Figure 1. (D) Histology of small intestine (left) and mesenteric AT (right) in Foxp3^RFP/GFP^ (WT, n = 2) and ΔtTreg (n = 3) mice on VegND, showing normal structure with no leukocytic infiltrates. Magnifications: small intestine, 200× (scale bar = 200 µm); mesenteric AT, 400× (scale bar = 40 µm). (E and F) Histology of VAT and SAT from 26-27-week-old Foxp3^RFP/GFP^ (WT), ΔtTreg, and ΔpTreg mice (n = 4). Magnifications: 200× (scale bar = 200 µm). (E) Mice fed HFD for 20 weeks, with infiltration scores (0 - 3) detailed in the Methods. (F) Age-matched mice on VegND showing no signs of leukocytic infiltrates. (G-J) Adipocyte quantification and metabolic tests in (G and H) VegND-fed and (I and J) HFD-fed mice (20 weeks). Symbols and error bars represent individual mice and mean ± SD. Unpaired t-test; p-values in Figure 1. Data in H are from one experiment representative of three independent experiments, data in J represent three independent experiments. (G and I) Adipocyte numbers and median cross-sectional areas were quantified from (E and F). (H and J) AUC analysis for ITT and GTT (VegND: n = 3 - 5mice/group; HFD: n = 6 - 10 mice/group).

Unsuspicious histopathological findings (*i.e.*, normal structure with no leukocytic infiltrates; Figure 2D, left)^29^ indicated that malnutrition due to intestinal inflammation can be excluded as the cause of the observed attenuated body weight gain of HFD-fed ΔtTreg mice. Compared to Foxp3^RFP/GFP^ mice, histologic analysis revealed largely comparable or slightly reduced immune infiltrates in both SAT and VAT of HFD-fed ΔtTreg mice, with similar adipose morphology, including adipocyte size and number (Figure 2E). When ΔtTreg mice were fed a VegND, no immune infiltration could be histologically detected in the mesenteric (Figure 2D, right), subcutaneous, and visceral (Figure 2F) AT. However, VegND-fed ΔtTreg mice exhibited a ’hypertrophic’ adipose morphology, as suggested by fewer but larger adipocytes per region of interest (Figure 2G). The latter correlated with increased insulin sensitivity and glucose tolerance (Figure 2H). These differences in adipocyte morphology were lost by HFD feeding (Figures 2E and I), whereas obesity-associated insulin resistance remained significantly attenuated in ΔtTreg mice even after 20 weeks of HFD feeding (Figure 2J).

### Selective pTreg cell paucity promotes obesity even with a normocaloric vegetarian diet

Next, we extended our studies to Foxp3^RFP/GFP^ mice selectively deficient in pTreg cells (ΔpTreg mice) due to a Foxp3-STOP allele that can be activated during intrathymic but not peripheral Treg cell development.^29,31^ Histological analyses revealed pronounced immune infiltrates in SAT and VAT of ΔpTreg mice when fed with HFD (Figure 2E), but not under VegND conditions (Figure 2F). However, adipocyte morphology in ΔpTreg mice remained largely unchanged, when compared to Foxp3^RFP/GFP^ mice (Figure 2G and I). Compared to ΔtTreg mice, the ablation of pTreg cells exacerbated obesity-associated insulin and glucose resistance, both under regular VegND (Figure 2H) and after 20 weeks of HFD feeding (Figure 2J) to a level that was comparable to Foxp3^RFP/GFP^ mice. In some experiments, however, ΔpTreg mice showed even poorer metabolic results than Foxp3^RFP/GFP^ mice, such as glucose tolerance test (GTT; Figure 2J).

In previous longitudinal studies in which VegND was fed, no significant differences in body weight were found between ΔtTreg and ΔpTreg mice up to 30 weeks of age.^29^ Here, we re-examined the body weight development of VegND-fed mice during age progression by extending the observation period up to 60 weeks. Compared to females, male wild-type mice are usually more prone to body weight gain, increased VAT inflammation, and the development of glucose intolerance and insulin resistance when fed with a HFD.^38–40^ However, even when fed with VegND, many ΔpTreg mice, both female (Figure 3A) and male (Figure 3B), developed a significantly higher body weight than ΔtTreg and Foxp3^RFP/GFP^ mice. While female ΔpTreg mice exhibited slightly earlier kinetics (Figure 3A), the excessive body weight gain in male ΔpTreg mice only became apparent from 20 weeks of age (Figure 3B). In fact, 44.1% of VegND-fed ΔpTreg males developed a body weight comparable to that of obese Foxp3^RFP/GFP^ mice after 20 weeks of HFD feeding (Figure 3B). The increased body weight of VegND-fed ΔpTreg mice correlated with marked increases in AT tissue weight (Figure 3C) and highly elevated levels of phosphorylated ERK (Figure 3D) associated with the development of overt insulin resistance and obesity.^41,42^ For comparison, in the AT of both ΔtTreg and Foxp3^RFP/GFP^ mice fed with VegND, levels of phosphorylated ERK protein were below the limit of detection, as expected (Figure 3D). Flow cytometric immunophenotyping of VegND-fed ΔpTreg mice indicated early signs of local immune dysregulation in ATs, in particular an increased compartment size of pro-inflammatory M1 macrophages at the expense of anti-inflammatory M2 macrophages (Figure 3E). Overall, these data provided the first evidence that even with physiologic caloric intake, selective pTreg cell deficiency predisposes ΔpTreg mice to spontaneously manifest impaired AT immune homeostasis and early signs of obesity.

**Figure 3.**
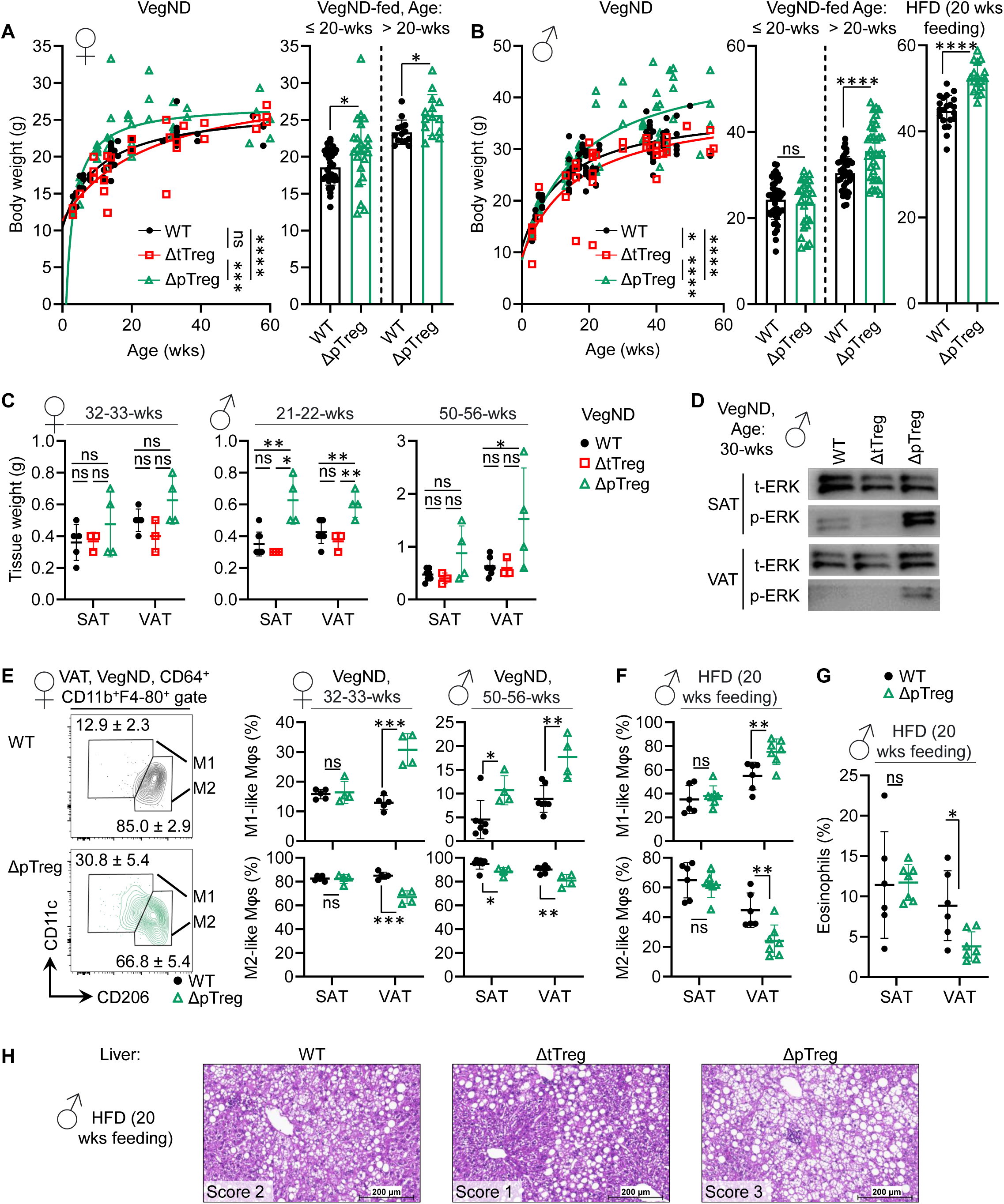
Obesity in ΔpTreg mice fed with a normocaloric vegetarian diet (A-E) Mice were fed a normocaloric VegND, unless otherwise noted. (G) Females - Body weight in ΔpTreg, Foxp3^RFP/GFP^ (WT) and ΔtTreg mice (n = 27 – 46) (H) Males - Left: Body weight in ΔpTreg, Foxp3^RFP/GFP^ (WT) and ΔtTreg mice (n = 39 – 86). Right: HFD-fed Foxp3^RFP/GFP^ (WT; n = 21) and ΔpTreg (n = 17) mice. (I) SAT and VAT weight in mice by sex and age, as indicated. (n = 3 - 8). (J) Western blot reveals elevated p-ERK levels in of SAT and VAT of ΔpTreg mice compared to Foxp3^RFP/GFP^ (WT) and ΔtTreg mice. (K) Shift to proinflammatory type 1 immunity in VAT of ΔtTreg mice. Representative contour plots (left) and cumulative M1 (top) and M2 (bottom) macrophage percentages in females (middle) and males (right) (n = 4 - 7). Mean percentages ± SD. (F-H) HFD feeding enhances shift to proinflammatory type 1 immunity in VAT of ΔpTreg mice. (L) Cumulative percentages of M1 (top) and M2 (bottom) macrophages in VAT. (M) Loss of eosinophils in VAT of ΔpTreg (n = 7) and Foxp3^RFP/GFP^ (WT; n = 6) mice. See also Figure S4. (N) Severe hepatic steatosis pronounced lipid accumulation in ΔpTreg mice, as revealed by liver histology from 26–27-week-old ΔpTreg (n = 2), Foxp3^RFP/GFP^ (WT; n = 3), and ΔtTreg (n = 3) mice. Symbols in (A, B) show individual mice. Symbols and error bars in (C, E, F, G) show individual mice and mean ± SD. Data in (F, G) from two experiments, representative of three independent experiments (n = 3 - 4 mice/experiment). Statistical analysis: non-linear regression for age vs. body weight (A, B); unpaired t-test (bar graphs in A, B; and C, E, F, G); p-values in Figure 1.

HFD feeding of ΔpTreg males further significantly enhanced the increase in body weight (Fig. 3B, right bar graph; Figure S4A), AT weight (Figure S4B), SVF cellularity (Figure S4C), and pro-inflammatory M1 macrophages (Figure 3F and S4D), while exacerbating the loss of EOs, compared to Foxp3^RFP/GFP^ mice (Figures 3G and S4D). The increased severity of obesity symptoms in the absence of pTreg cells was also illustrated by pronounced hepatic steatosis of HFD-fed ΔpTreg mice, including massive immune infiltrates and lipid deposition (Figure 3H).

### CD4^+^ T cell responses in the VAT of ΔtTreg and ΔpTreg mice

So far, our data suggest that tTreg and pTreg cells exert opposing functions in VAT homeostasis and obesity, as evidenced by the analysis of Foxp3^RFP/GFP^ mice and their tTreg and pTreg cell-deficient derivatives. This was reflected, among other things, in alterations of innate VAT immune cell compartments in ΔtTreg (Figure S3C and D) and ΔpTreg (Figures 3E-G and S4D) mice. We observed no quantitative differences between T conventional cells in VAT of ΔtTreg and ΔpTreg mice (Figures 4A and S5A). However, we noticed qualitative differences, *e.g.* in the percentages of CD4^+^ T cells with a naïve (CD62L^high^CD44^low^) and an effector/memory (CD62L^low^CD44^high^) phenotype (Figures 4B and S5B). To gain mechanistic insights into Teff cell responses, FACS-purified CD4^+^ T cells from the VAT of mice fed HFD for 20 weeks (Figure 4C) were subjected to single-cell transcriptomics. Based on differential expression of canonical marker genes (Figure S6, Table S1), the VAT CD4^+^ T cells of all three HFD-fed mouse lines (Foxp3^RFP/GFP^, ΔtTreg, and ΔpTreg) were grouped into eight distinct clusters of cells with similar gene expression patterns (Figure 4D). As revealed by Uniform Manifold Approximation and Projection (UMAP) visualization, a common feature was the predominance of cell clusters with pro-inflammatory Th1 properties, notably the ’Th1’ cluster defined by high mRNA expression of genes with well-known functions in Th1 cells (*e.g.*, *Tbx21*, *Ly6c2*, *Nkg7*, *Ifng, Fasl*, *Ctsw*, *Tnf*, *Cxcr3,* and *Ccl5*) (Figures 4D and S6; Table S1). The other Th1-like clusters (Figures 4D and E) further expressed high mRNA levels of markers indicating a more advanced stage of Teff cell differentiation (Th1/Tex; Tex, Tmem, and CTL). This included putative markers of T cell exhaustion and mediators of chronic VAT inflammation (’Th1/Tex’ cluster: *Tbx21, Nkg7, Ifng, Pdcd1, Lag3, Tnfsf8, Il21*, *Spp1*, and *Eea1*; ’Tex’ cluster: *Lag3, Tnfsf8, Il21*, *Spp1*, *Eea1, Sostdc1, Cd200, Izumo1r, Slamf6, Gpm6b*, and *Tbc1d4*). In addition to its resemblance to Th1 cells, the ’CTL’ cluster was characterized by high expression levels of *Eomes* and *Gzmk*, while the ’Tmem’ cluster exhibited a downregulated naïve T cell signature (*Sell*, *Lef1*, *Ccr7*, *Foxp1*, *Igfbp4*, *Satb1,* and *Dapl1*) with concomitantly upregulated expression of effector/memory-type markers (*e.g*., *Cd44* and *Il7r*) (Figures 4D and S6; Table S1).

**Figure 4.**
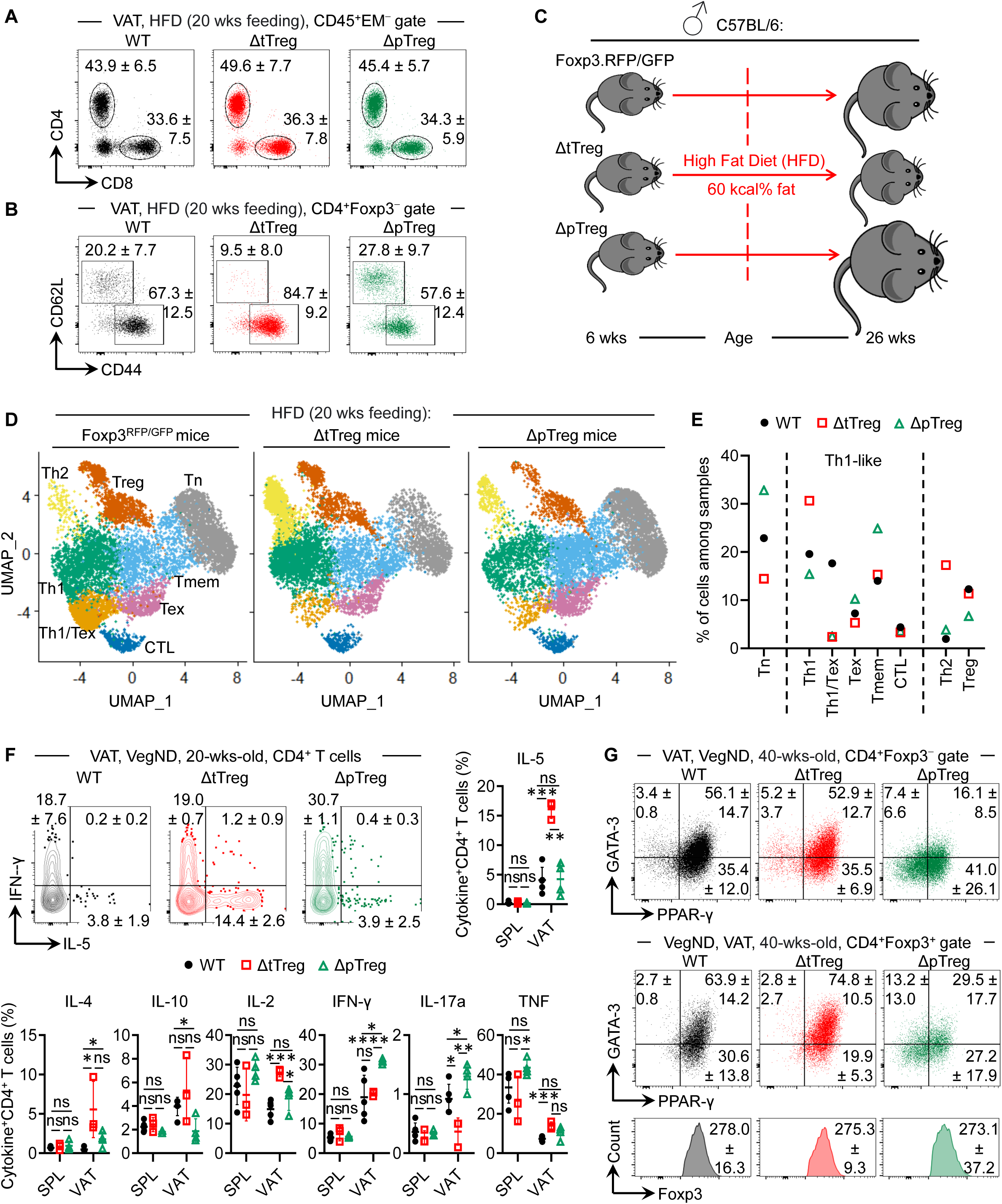
CD4^+^ T cell responses in the VAT of ΔtTreg and ΔpTreg mice (A-B) Flow cytometry of T cells in VAT of HFD-fed Foxp3^RFP/GFP^, ΔtTreg, and ΔpTreg mice. Data represent three independent experiments (n = 6 - 11 mice/group). See Figures S5A and S5B. (A) Representative dot plots show frequency of CD4^+^ and CD8^+^ T cells among gated CD45^+^EM^−^ cells. (B) Activation status of gated CD4^+^Foxp3^−^ cells as revealed by CD44 and CD62L expression. (C-E) scRNA/TCR-seq of CD4^+^ T cells from VAT of HFD-fed mice. (C) Experimental design. After 20 weeks of HFD feeding, CD4^+^ T cells were isolated from VAT of Foxp3^RFP/GFP^, ΔtTreg, and ΔpTreg mice for scRNA/TCRseq. See Figure S11. (D) UMAP depicting eight distinct cell clusters across individual samples. See Figure S6 for expression of key markers used for cluster definition. (E) Size of cell clusters shown in (D) across Foxp3^RFP/GFP^, ΔtTreg, and ΔpTreg mice. (F-G) Flow cytometry of CD4^+^ T cells in VegND-fed Foxp3^RFP/GFP^, ΔtTreg, and ΔpTreg mice. (F) Cytokine production after PMA/ionomycin restimulation *in vitro*. Representative flow cytometry and cumulative percentages of cytokine^+^CD4^+^ T cells from VAT and spleen (SPL), as indicated. (G) Intracellular staining of GATA-3 and PPAR-γ in CD4^+^Foxp3^−^ Tcon (top) and CD4^+^Foxp3^+^ Treg (middle) cells, and Foxp3 expression (bottom) in VAT. Mean percentages or MFI ± SD. (n = 3 - 5 mice/group).

Relevant to the mechanisms underlying divergent disease manifestation in HFD-fed ΔtTreg and ΔpTreg mice, UMAP visualization allowed several important conclusions to be drawn: compared to Foxp3^RFP/GFP^ and ΔpTreg mice, attenuated pro-inflammatory type 1 immune responses and obesity symptoms correlated with enhanced anti-inflammatory Th2 CD4^+^ T cell responses of ΔtTreg mice, as indicated by a markedly increased cell cluster with a Th2 phenotype (’Th2’ cluster: *Gata3*, *Hilpda*, *Nfkbia*, *Tnfsf11*, *Atf3*, *Ltb4r1*, *Ramp3*, *Ccr2*, *and Il1rl1*) (Figures 4D, 4E; and S6; Table S1). The concomitant reduction of the naïve T cell (’Tn’) cluster (*Sell*, *Lef1*, *Ccr7*, *Foxp1*, *Igfbp4*, *Satb1,* and *Dapl1*) (Figures 4D and 4E) was consistent with our flow cytometric analyses (Figures 4B and S5B), suggesting a scenario of enhanced naïve CD4^+^ T cell differentiation into Th2-like cells in VAT of HFD-fed ΔtTreg mice, although we cannot formally exclude impaired local recruitment of naïve T cells by migration and homing into VAT. Whereas, the increased ’Th1’ cluster and the concomitantly decreased ’Th1/Tex’ cluster of HFD-fed ΔtTreg mice provided evidence for a blunted Th1 response (*e.g.*, due to abrogated progression to an advanced pathogenic Th1 effector cell stage).

In HFD-fed ΔpTreg mice, an exacerbated pro-inflammatory type 1 immune response and obesity symptoms correlated with a reduced ’Treg’ cluster (*Foxp3*, *Il1rl1*, *Il2ra*, *Ikzf2*, *Ctla4*, *Tnfrsf9*, *Klrg1*, and *Areg*) (Figures 4D and 4E), whereas the size of the ’Tn’ and ’Tmem’ clusters were increased (Figures 4D and 4E) (See Figure 4B for comparison). Even when fed normocaloric VegND, the proportion of VAT CD4^+^ T cells producing the pro-inflammatory cytokines IFN-γ (Figure 4F) and IL-17 (Figure S5C), as revealed by intracellular staining after a brief T cell restimulation *in vitro*, was significantly increased in ΔpTreg mice. In contrast, the VAT of VegND-fed ΔtTreg mice was characterized by significant increases in the proportion of CD4^+^ T cells producing IL-2 (Treg cell growth factor) and anti-inflammatory Th2 cytokines, including IL-4, IL-5, and IL-10 (Figures 4F and S5C). Consistently, both CD4^+^ T conventional and Foxp3^+^ Treg cells in the VAT of ΔtTreg mice exhibited a predominantly Th2- like PPAR-y^+^GATA-3^+^ phenotype, similar to that observed in Foxp3^RFP/GFP^ mice (Figure 4G). With the exception of increased IL-10, we did not observe any significant differences in cytokine expression among SAT CD4^+^ T cells in ΔtTreg compared to Foxp3^RFP/GFP^ mice (data not shown).

In summary, our single-cell transcriptomics data from HFD-fed mice show distinct qualitative differences in VAT-CD4^+^ T cell responses that correspond well with their divergent manifestation of disease symptoms. Unexpectedly, this anti-inflammatory Th2 bias in ΔtTreg mice and the pro-inflammatory Th1 bias in ΔpTreg mice can already be observed, at least in part, in mice with physiological calorie intake.

### VAT pTreg cells resemble tissue-type Treg cells

In the VAT of ΔtTreg mice, pTreg cells numerically compensated for selective tTreg cell deficiency, as indicated by the similar size of the UMAP ’Treg’ cluster (Figures 4D and 4E) and the increased RFP^+^GFP^−^ pTreg cell compartment (Figures 5A and 5B) compared to that of Foxp3^RFP/GFP^ mice. The VAT Treg cells of both ΔtTreg and Foxp3^RFP/GFP^ mice formed multiple micro clones (≥ 2, ≤ 21 cells), with a comparable frequency of clonally expanded cells (Foxp3^RFP/GFP^: 38.9%; ΔtTreg: 41.9%), although shared TCR clonotypes between different samples were not detected (Figure 5C). Overall, this finding is consistent with previous studies showing that VAT Treg cells can form multiple micro clones in individual samples, while maintaining a high degree of diversity between different mice.^7,27^ Our scRNA-seq data revealed that tTreg cell ablation further enhanced the expression of many genes of the so-called ’VAT-Treg cell signature’^10,11,14,21,27^ in the VAT ’Treg’ cluster of ΔtTreg mice (Figure 5D). This included genes encoding transcription factors that promote AT adaptation (*e.g.*, PPAR-γ and GATA-3), chemokine receptors mediating VAT homing (*e.g.*, CXCR6, CCR1, and CCR3), and tissue-type effector molecules (*e.g.*, ST-2, KLRG1, CD127, and IL9R), including T cell activation markers (*e.g.*, CD44) and molecules involved in lipid metabolism (*e.g.*, DGAT1) (Figure 5D). The expression levels of many genes encoding proteins with well-known functions in Treg cell-mediated suppression [*e.g.*, TGFβ, Fibrinogen-like protein 2 (FGL2), IL10RA, CD40L, Galectin-1, and TRAIL]^43–45^ were also upregulated in the VAT ’Treg’ cluster of ΔtTreg mice, as compared to Foxp3^RFP/GFP^ mice (Figure 5E). Consistently abrogated in the VAT ’Treg’ cluster of ΔtTreg mice was also the expression of a more comprehensive set of genes (Figure 5F; ’obese VAT’ Treg cell signature) known to be either upregulated or downregulated in total VAT Treg cells in HFD-induced obesity.^15^

**Figure 5.**
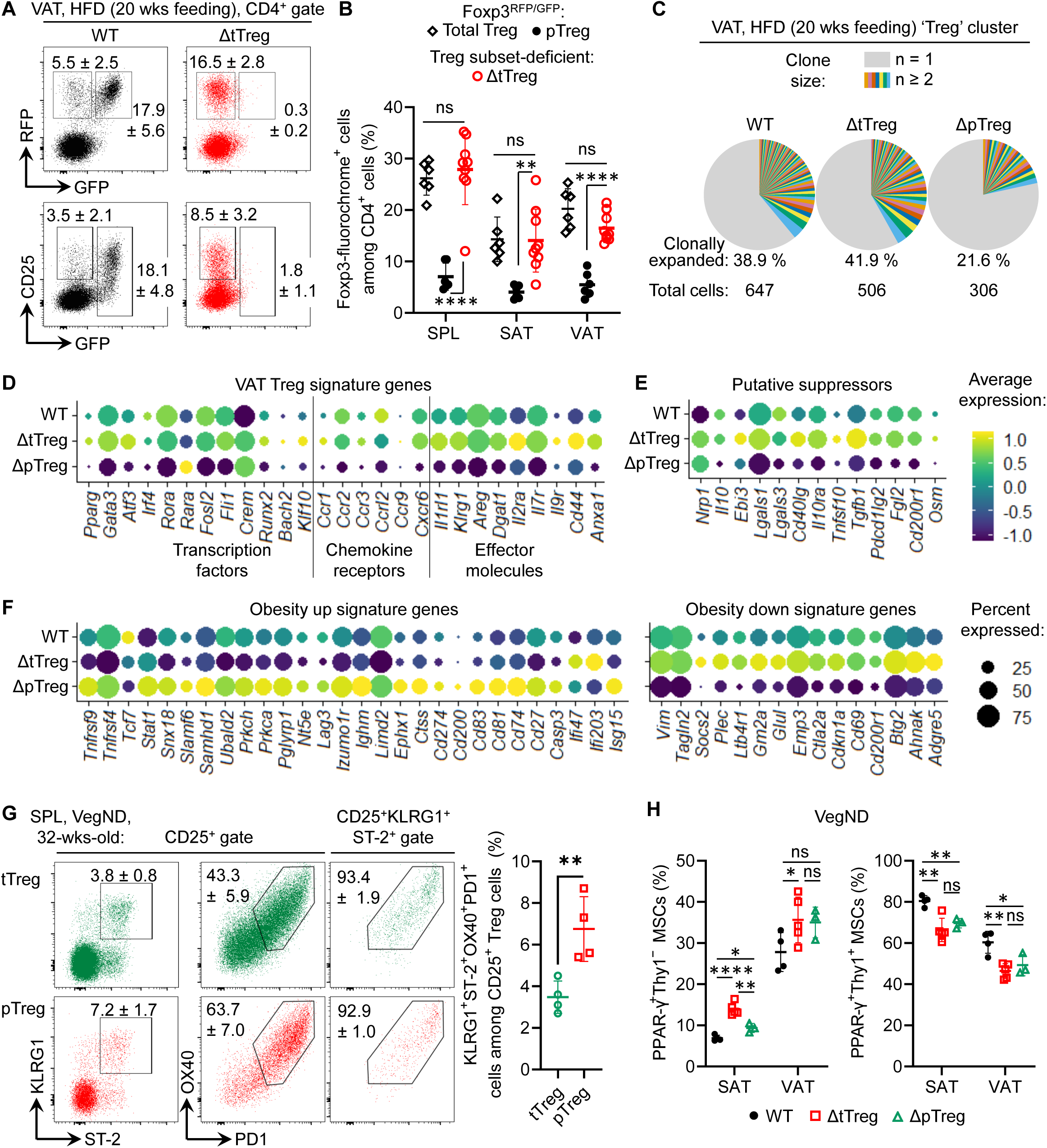
pTreg cells resemble tissue-type Treg cells (A-B) pTreg cells compensate for tTreg cell loss in HFD-fed ΔtTreg mice. (A) Flow cytometry of Foxp3-driven RFP vs. GFP (top) and CD25 vs. RFP expression (bottom) in CD4-gated cells from VAT of Foxp3^RFP/GFP^ (WT; n = 6) and ΔtTreg (n = 9) mice. Data are from two experiments, representative of nine independent experiments (≥ 3 mice/group). For comparability, WT data shown here are also presented in Figures 6A and 6B. (B) Cumulative percentages of total Treg and pTreg cells from (A) in Foxp3^RFP/GFP^ and ΔtTreg mice. (C-F) Comparative analysis of the ’Treg’ cluster across Foxp3^RFP/GFP^ (WT), ΔtTreg, and ΔpTreg mice using scRNA/TCRseq data from Figure 4D. (C) Pie charts depicting TCR clone frequencies, categorized by clone size (1 vs. ≥2). (D) VAT Treg signature genes. (E) Genes implicated in suppression. (F) Genes either upregulated (left) or downregulated (right) in Treg cells during obesity. (G) Representative flow cytometry and cumulative percentages of tissue-type Treg precursors among splenic tTreg (top) and pTreg (bottom) cells in 32-week-old Foxp3^RFP/GFP^ males fed VegND. (H) Flow cytometry of adipogenic MSCs in VegND-fed Foxp3^RFP/GFP^ (WT), ΔtTreg, and ΔpTreg females. Composite percentages of PPAR-γ^+^ MSC subsets that are either Thy1^−^ (left) or Thy1^+^ (right) among gated CD45^+^CD31^+^Pdgfrα^+^Sca1^+^ cells. (n = 3 -5 mice/group). See also Figure S7. Numbers in (A, G) represent mean percentages ± SD. Data in (G, H) are from a single experiment (n = 4), representative of two to three independent experiments. Unpaired t-test (B, G, H); p-values in Figure 1.

We conclude that in HFD-fed ΔtTreg mice, the up to 5-fold proportional increase in VAT pTreg cells (Figure 5B) and their phenotypic adaptations (Figures 5D-F), including sustained expression of genes typically downregulated in obesity (‘obesity-down signature’), counteract the transcriptional Treg aberrations linked to chronic VAT inflammation. These data attribute a key functional role to pTreg cells in VAT homeostasis, which was not expected given their small contribution to the physiological VAT Treg cell pool.

### Contribution of pTreg and tTreg cell development to tissue-type Treg cell precursors

Tissue-type Treg cells present in non-lymphoid tissues that express ST-2 have been shown to arise from splenic Foxp3^+^ Treg cell precursors and to accumulate in non-lymphoid tissues, where they undergo further adaptations to fulfil homeostatic and regenerative functions.^21^ Consistently, based on physiologic frequencies of clonally expanded cells (Figure 5C), the markedly increased population size of pTreg cells in VAT of ΔtTreg mice (Figures 5A and 5B) is likely to be driven by enhanced recruitment, rather than clonal expansion of pre-formed VAT pTreg cells. Previous transcriptomics analyses using Foxp3^RFP/GFP^ mice have already indicated striking similarities between the core transcriptional signature of pTreg cells (but not tTreg cells)^25^ and that of tissue-resident Treg cells at steady-state,^21,46,47^ which included increased expression levels of ST-2 and KLRG1.^25^ Here, we utilized Foxp3^RFP/GFP^ mice to directly track the developmental origin tissue-type Treg cell precursors, as defined by expression of CD25, KLRG1, ST-2, OX40, and PD-1.^21^ The results show that such tissue-type Treg cell precursors can originate from both thymic and peripheral Treg cell development (Figure 5G), but were significantly enriched in the pTreg (6.8 ± 1.6%) as compared to the tTreg cell compartment (3.5 ± 0.8%). Consistent with a key role of VAT Treg cells in restraining the differentiation of PPAR-γ^+^ mesenchymal stromal cells (MSCs) into adipocytes through oncostatin M-dependent mechanisms,^13^ we found *Osm* mRNA levels to be selectively increased in the ’Treg’ cluster of ΔtTreg mice (Figure 5E). In ΔtTreg mice, we further found that the size of one of the subsets proposed to have adipogenic potential, PPAR-γ^+^Thy1.1^−^ MSCs,^13,48^ was significantly increased (Figures 5H and S7). This accumulation of PPAR-γ^+^Thy1.1^−^ MSCs is in line with the attenuation of their pTreg cell-mediated developmental progression and the ’hypertrophic’ adipose morphology observed in ΔtTreg mice (Figures 2F and G). Such Thy1.1^−^MSCs can give rise to another progenitor population, Thy1.1^+^ MSCs,^13,48^ which were markedly reduced in ΔtTreg mice (Figure 5H). This is consistent with continued development of Thy1.1^+^ MSCs into adipocytes while their *de novo* formation from Thy1.1^−^ MSCs was blocked by increased pTreg cell activity.

### Numerical and functional impairment of Treg cells in VAT of ΔpTreg mice

Our observation that the Treg cell reduction in VAT of HFD-fed Foxp3^RFP/GFP^ mice is due to selective loss of the tTreg cell subset (Figures 1H, 1I, and S2B) raised the question of how HFD impinges on the maintenance of the VAT Treg cell pool in ΔpTreg mice, which consists exclusively of tTreg cells, precluding pTreg cell-mediated compensatory mechanisms. In fact, in HFD-fed ΔpTreg mice, the size of the VAT Treg cell pool remained well below that of Foxp3^RFP/GFP^ mice (∼50% reduction) (Figure 6A). This was consistent with a substantially reduced ’Treg’ cluster size (∼46% reduction) as revealed by UMAP visualization (Figures 4D and 4E) and frequency of clonally expanded VAT Treg cells (∼50% reduction) (Figure 5C), indicating that VAT tTreg cells of ΔpTreg mice undergo limited proliferative expansion *in situ*. A reduction in the size of the Treg cell compartment was also observed in SAT and peripheral lymphoid tissues (Figure 6B), although less pronounced than in VAT of ΔpTreg mice.

**Figure 6.**
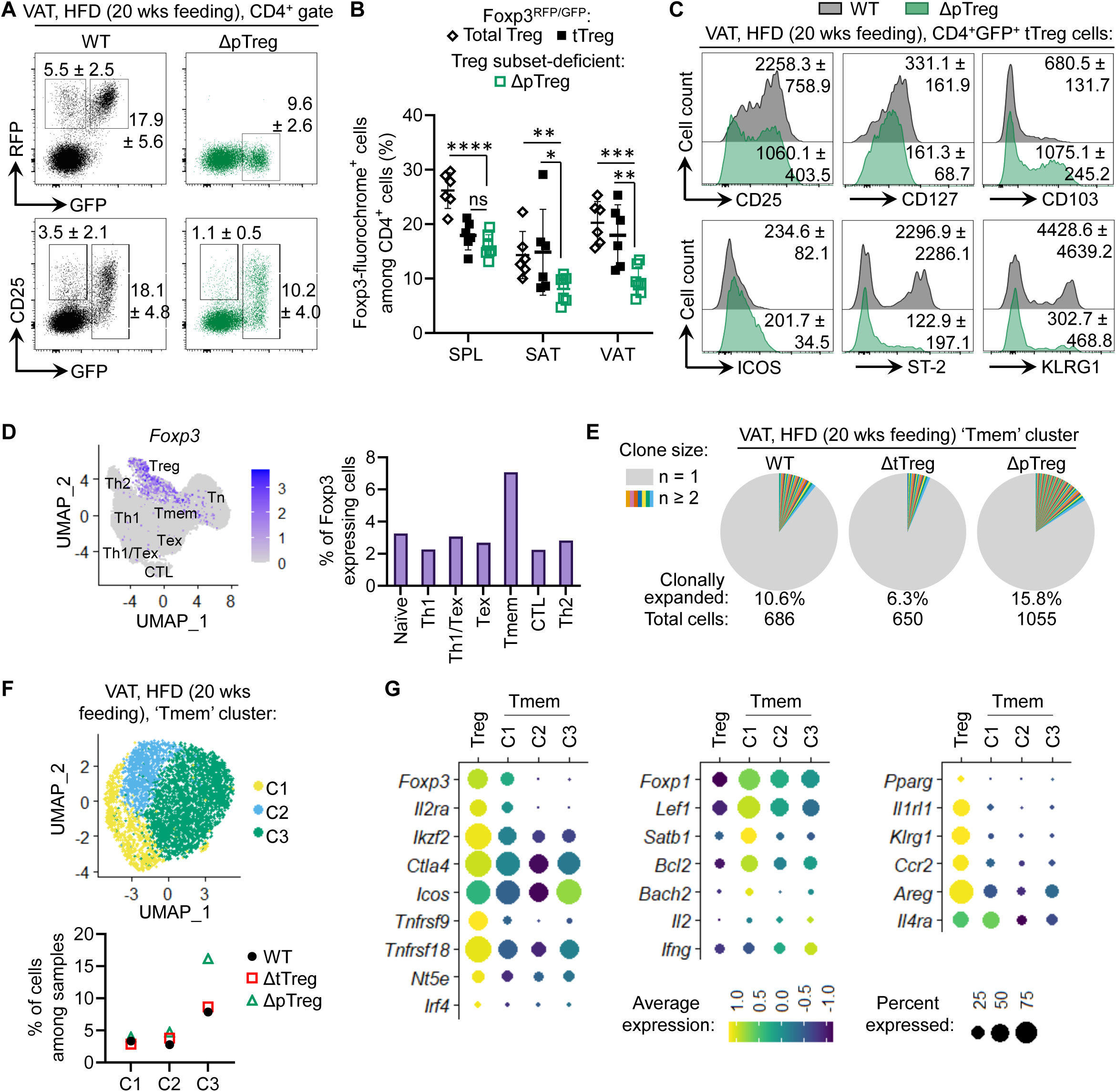
Numerical and functional impairment of VAT Treg cells in HFD-fed ΔpTreg mice (A-C) Flow cytometry of Treg cells in HFD-fed Foxp3^RFP/GFP^ (n = 6) and ΔpTreg (n = 7) mice. (A) Flow cytometry of Foxp3-driven RFP vs. GFP (top) and CD25 vs. RFP expression (bottom) in CD4-gated VAT cells from VAT of Foxp3^RFP/GFP^ (WT) and ΔpTreg mice. Mean ± SD. Data are from two experiments, representative of nine independent experiments (≥ 3 mice/group). For comparability, WT data shown here are also presented in Figures 5A and 5B. (B) Cumulative percentages of total Treg and tTreg cells from (A) in Foxp3^RFP/GFP^ and ΔpTreg mice. Individual mice and mean values ± SD. Unpaired t-test; p-values in Figure 1. (C) Dysregulated VAT tTreg cell signature of ΔpTreg mice. Histograms show selected surface markers in VAT tTreg cells (CD4^+^GFP^+^) of WT (grey) and ΔpTreg (green) mice. Mean MFI ± SD. See also Figure S8. Data from two experiments, representative of at least three independent experiments (≥ 3 mice/group). (D) UMAP visualization of relative *Foxp3* mRNA expression in VAT CD4^+^ T cells (left) and frequency of *Foxp3* mRNA^+^ cells in non-Treg clusters in VAT (right) based on scRNA-seq data depicted in Figure 4D. (E-G) Comparative analysis of the ’Tmem’ cluster across Foxp3^RFP/GFP^ (WT), ΔtTreg, and ΔpTreg mice using scRNA/TCRseq data from Figure 4D. (E) Enhanced clonal expansion in the ’Tmem’ cluster of ΔpTreg mice. Pie charts depict TCR clone frequencies in the VAT ’Tmem’ cluster, categorized by clone size (1 vs. ≥2). (F) Sub-clustering of the ’Tmem’ cluster. UMAP visualization of the VAT ’Tmem’ cluster (top) and size of sub- clusters among VAT CD4^+^ T cells (bottom). (G) Selected gene expression across the ’Treg’ cluster and ’Tmem’ sub-clusters from (F), highlighting Foxp3- stabilized (left), Foxp3-repressed (middle), and VAT Treg signature genes (right). See also Figure S9.

In residual VAT Treg cells, the expression levels of the Foxp3-fluorochrome reporter (Figure 6A) and Foxp3 protein (Figure 4G) was comparable to that of Foxp3^RFP/GFP^ mice. In contrast, a selected set of proteins of the ’VAT Treg cell’ signature was down-regulated in VAT Treg cells of ΔpTreg mice, including key transcription factors for VAT adaptation (*e.g.*, PPAR-γ, GATA-3) (Figure 4G) and functionally relevant surface markers (*e.g.*, CD25, CD127, ST-2, and KLRG1) (Figure 6C and S8), showing a high correlation between protein and mRNA expression (Figure 5D). Overall, selective pTreg cell ablation generated pronounced ’holes’ in the transcriptional ’VAT Treg cell’ signature (Figure 5D) that extended to genes encoding proteins with known functions in suppressor mechanisms (Figure 5E). Conversely, the ’obese VAT’ Treg cell signature whose expression correlates with HFD-induced obesity in Foxp3^RFP/GFP^ mice was consistently enhanced in VAT Treg cells of ΔpTreg mice (Figure 5F), reflecting their exacerbated obesity symptoms (Figures 3 and S4).

### Treg cell destabilization and dedifferentiation into T effector cells in VAT of ΔpTreg mice

Because many ’VAT-Treg cell’ signature genes are under direct Foxp3 transcriptional control,^49,50^ we re-examined our single-cell transcriptomics data. We paid particular attention to cells in the seven ’non-Treg’ clusters, which have transcriptional similarities to the ’Treg’ cell cluster but were not considered Treg cells in our original cluster analysis due to low *Foxp3* mRNA expression. Outside the ’Treg’ cluster, we found a significant number of CD4^+^ T cells that express low but clearly detectable levels of *Foxp3* mRNA, which were particularly enriched in the ’Tmem’ cluster of ΔpTreg mice (Figure 6D). The ’Tmem’ cluster of ΔpTreg mice was further characterized by an increase in size (Figure 4D and 4E) and an accumulation of clonally expanded cells (Figure 6E), and could be subdivided into three sub-clusters (C1-C3) (Figure 6F). The ’Tmem’ sub-cluster C1 was enriched for cells expressing low levels of *Foxp3* mRNA and other ’Treg’ signature genes (*e.g.*, *Il2ra*, *Ikfz2*, *Il4ra*, and *Cd74*) (Figure S9; Table S2). Consistent with low levels of *Foxp3* mRNA, all ’Tmem’ sub-clusters exhibited reduced mRNA levels (Figure 6G, left) of genes whose expression is enhanced and stabilized by Foxp3 (*e.g.*, *Foxp3* itself, *Il2ra*, *Ikzf2*, *Tnfrsf9*, and *Ctla4*).^50,51^ Conversely, the expression of genes repressed by Foxp3^50,51^ were up-regulated (*e.g.*, *Foxp1*, *Lef1*, *Bcl2*, and *Il2*) (Figure 6G, middle). Additionally, genes of the ’VAT-Treg cell’ signature were consistently downregulated (Figure 6G, right). Overall, the gradual loss of *Foxp3*, *Il2ra, Helios,* and *Ctla4* mRNA expression, as well as the gradual increase in *Ifng* and *Il2* mRNA expression (*i.e.*, ’Treg’ > C1 > C2 > C3) suggested that ’Tmem’ sub-cluster C1 harbors an unstable Treg cell population, which is prone to differentiate into ’Tmem’ sub-cluster C3. In contrast to C1 and C2, which exhibited a more naïve-like phenotype (e.g., *Sell*), ’Tmem’ sub-cluster C3 was characterized by a Teff-like phenotype expressing genes (*e.g*., *Tnfrsf4*, *Tnfsf8*, *Pdcd1*, *Nr4a3*, and *Tigit*), which have been implicated in the pathogenesis of VAT inflammation^52–54^ (Figure S9; Table S2). Notably, Tmem’ sub- cluster C3 increased in size in ΔpTreg mice, compared to Foxp3^RFP/GFP^ and ΔtTreg mice (Figure 6F). In conclusion, the numerical and functional impairment of VAT Treg cell pool in ΔpTreg mice can be attributed by the loss of Treg cell identity and their dedifferentiation into ’ex’-Treg cells with an T cell effector/memory phenotype.

### Correlating the accumulation of ex-tTreg cells in VAT with severity of obesity symptoms

Dedifferentiation of VAT tTreg cells into pathogenic Foxp3^−^ Teff cells in HFD-fed ΔpTreg mice (Figure 6) provides both a mechanistic explanation for the selective loss of tTreg cells in VAT of HFD-fed Foxp3^RFP/GFP^ mice (Figures 1H, 1I, and S2B) and for the attenuation of clinical symptoms, observed in HFD-fed ΔtTreg mice (Figures 2 and S3). To directly track and quantify such ex-Treg cells, we performed genetic fate-mapping studies in R26Y x Foxp3^RFP/GFP^ mice, which additionally express YFP in all tTreg lineage cells, regardless of their Foxp3 protein expression status (Figures 7A and 7B). This allows to track RFP^−^YFP^+^ ex-tTreg cells despite their loss of Foxp3-dependent reporter expression (Figure 7B).^31^ Compared to ND-fed mice, 10 weeks of HFD feeding significantly increased the accumulation of ex-Treg cells in VAT (Figure 7C and D), with up to 50% of VAT CD4^+^ T cells exhibiting an RFP^−^YFP^+^ ex-Treg cell phenotype (Figure 7C; mouse #2).

**Figure 7.**
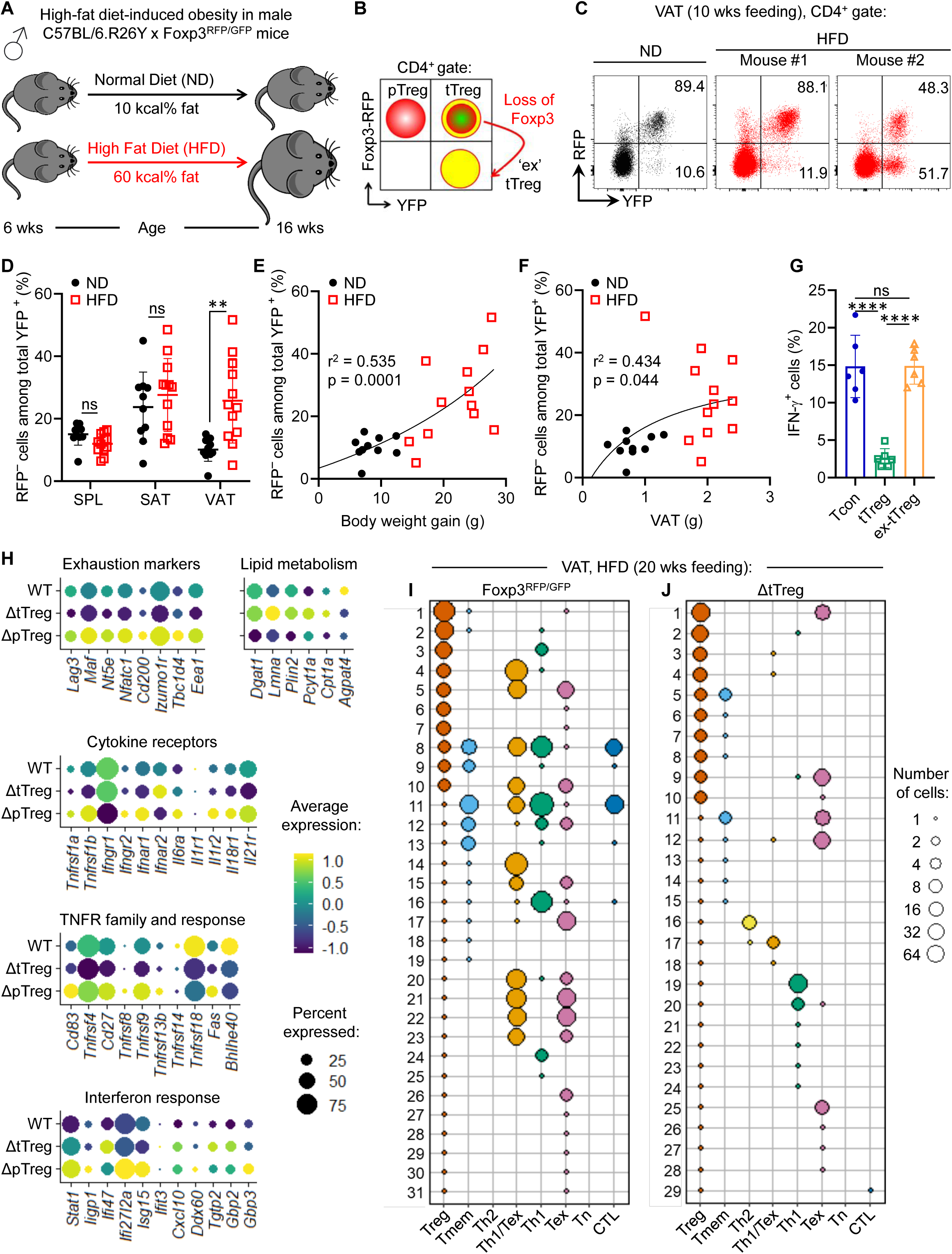
Correlation of ex-tTreg cell accumulation in VAT with obesity severity (A) R26Y x Foxp3^RFP/GFP^ mice were fed HFD for 10 weeks, with ND-fed mice as controls. (B) YFP-based lineage tracing of tTreg cells in R26Y x Foxp3^RFP/GFP^ mice. (C-F) Flow cytometry of HFD-induced VAT tTreg cell dedifferentiation. (C) Representative dot plots of RFP vs. YFP expression among CD4-gated cells in VAT. Percentages among total YFP^+^ cells. (D) Cumulative percentages of RFP^−^ ex-tTreg cells among total YFP^+^ cells. (E and F) Correlation of VAT tTreg cell dedifferentiation with (E) body weight gain and (F) VAT weight (n = 10 - 12 mice/group). Pearson correlation analysis with r^2^ as coefficient of determination. Data represent three independent experiments (n = 3 - 4 mice/experiment). (G) IFN-γ production by VAT ex-tTreg cells. Frequency of IFN-γ^+^ VAT Tcon, tTreg, and ex-tTreg cells from HFD- fed R26Y x Foxp3^RFP/GFP^ mice (20 weeks) after *in vitro* PMA/Ionomycin stimulation. Data from one experiment (n = 6). See also Figure S10. (H) Gene expression in the ’Treg’ cluster from Figure 4D. Bubble plots show average and percent expression of selected genes in Foxp3^RFP/GFP^ (WT), ΔtTreg, and ΔpTreg mice. (I and J) Treg cell clonotype distribution. VAT Treg clonotype (rows) across CD4^+^ T cell clusters from Figure 4D in (I) Foxp3^RFP/GFP^ and (J) ΔtTreg mice, based on CDR3 amino acid sequences of TCR-α and -β chains. Unpaired t-test (D, G); p-values in Figure 1. Symbols in (D-G) represent individual mice; bars show mean ± SD.

Several observations ruled out the possibility that such Foxp3^−^YFP^+^ CD4^+^ T cells in VAT arose from transient intrathymic Foxp3 expression during conventional T cell development,^55,56^ rather than from local dedifferentiation of VAT Foxp3^+^ tTreg cells into Foxp3^−^ ex-tTreg cells: First, and consistent with local destabilization and dedifferentiation of VAT tTreg cells, flow cytometry (MFI) of VAT RFP^+^YFP^+^ tTreg cells and Foxp3^−^YFP^+^ cells indicated a gradual loss of proteins (Figure S10A) encoded by genes under direct Foxp3 transcriptional control, including Foxp3 itself, CD25, and ST-2. In fact, reduced expression levels of VAT tTreg cell surface markers closely correlated with the frequencies of Foxp3^−^YFP^+^ cells in VAT of HFD-fed mice (Figure S10B). Second, FACS-isolated Foxp3^−^YFP^+^ cells from VAT of HFD-fed mice (Figure S10C) rapidly reacquired Foxp3 expression upon stimulation *in vitro* (Figure S10D), a characteristic of ex-Treg cells distinguishing them from thymus-derived Foxp3^−^YFP^+^ cells that transiently expressed Foxp3.^55,56^

Overall, the extent of tTreg cell dedifferentiation and ex-tTreg cell accumulation differed between individual HFD-fed R26Y x Foxp3^RFP/GFP^ mice and correlated with their gain in body weight (Figure 7E) and VAT weight (Figure 7F). Furthermore, the profile of effector cytokines produced by FACS-isolated VAT ex-tTreg cells upon brief re-stimulation *in vitro* (Figures 7G and S10E) was predominated by IFN-γ (Figure 7G), while only a few cells expressed low amounts of IL-2 (data not shown). This correlation between VAT tTreg cell dedifferentiation and clinical signs of obesity further indicated that the resultant ex-tTreg cells indeed feed into the pool of pathogenic Th1 cells, thereby fueling HFD-induced type 1 inflammation.

Mechanistically, differential expression of mRNAs implicated in T cell exhaustion, lipid metabolism, and cytokine signalling (Figure 7H) pointed towards both metabolic^11,57,58^ and inflammatory^15,59,60^ cues that locally destabilize tTreg cells in the obese VAT (Figure 7C) but not SAT or spleen (Figure 7D). In particular, the UMAP ’Treg’ cluster of ΔpTreg mice was characterized by high expression levels of mRNAs encoding several receptors for pro-inflammatory cytokines well-known for their Treg cell-destabilizing activity, including TNF, IFN-α, and IL-6 (Figure 7H). The functional relevance of increased pro-inflammatory cytokine receptor expression was corroborated by pathway analyses, revealing high expression levels of mRNAs encoding TNF and IFN receptor response genes in VAT tTreg cells of obese ΔpTreg mice (Figure 7H, bottom).

### Direct clonal relationship between tTreg cells and pro-inflammatory Foxp3^−^ T effector cells

Further efforts to evaluate the pathogenic potential of such ex-tTreg cells in functional assays, such as adoptive T cell transfer, were hampered by the small number of cells we were able to isolate by FACS from the VAT of HFD-fed mice. However, we did analyze the developmental trajectory and clonal relationship between Foxp3^+^ Treg and Foxp3^−^ Teff cells in the obese VAT. For this, the TCR clonotypes of the ’Treg’ cluster were traced within the other VAT CD4^+^ T cell clusters obtained from our scRNA/TCR-seq data. Our analysis revealed that the ’Treg’ cluster of Foxp3^RFP/GFP^ mice shared many clonotypes with the pro-inflammatory ’Th1’ cluster (*i.e.*, ’Th1’, ’Th1/Tex’, ’Tex’, and ’Tmem’) (Figure 7I).

The majority of such shared clonotypes were unique within the ’Treg’ cluster (n = 1), but clonally expanded among the pathogenic Teff cell clusters (n ≥ 2). Notably, many of the unique ’Treg’ cluster clonotypes were recovered from the Teff cell clusters at a rather high frequency of up to 60 cells (Figure 7I). This indicates that VAT tTreg cells continue to undergo proliferative expansion after local antigen recognition via their TCRs and loss of Foxp3 expression in the highly pro-inflammatory VAT microenvironment of HFD-fed Foxp3^RFP/GFP^ mice. As expected, we did not observe any such Treg cell clonotypes in the ’Tn’ and ’Th2’ clusters, which also rules out mRNA contamination between individual cells as. a major source of the repeatedly identified TCR clonotypes. Lastly, genetic tTreg cell ablation greatly reduced the accumulation of ’Treg’ cluster clonotypes in the ’non-Treg’ clusters (Figure 7J). This finding further supports the conclusion that tTreg cells are the main population contributing to the shared clonotypes detected in VAT of HFD-fed Foxp3^RFP/GFP^ mice.

## DISCUSSION

With the overarching aim of unraveling the significance of VAT Treg cell developmental heterogeneity, we have identified pTreg cells as pivotal regulators of VAT homeostasis, while tTreg cells play a critical role in promoting obesity. A key strength of our experiments is the use of the dual Foxp3^RFP/GFP^ mouse model, along with its ΔpTreg and ΔtTreg derivatives, which allowed for comprehensive studies both under physiological conditions and in the absence of pTreg or tTreg cells. This approach not only allowed for the dissection of their individual contributions, but also controlled for potential confounding variables, and provided insights into compensatory mechanisms. Such studies have been previously hindered by the lack of suitable loss-of-function models. Foxp3.CNS1^−/−^ mice^61^ exhibit a selective reduction of pTreg cells, but express an N-terminal Foxp3-GFP fusion protein that acts as a hypomorph,^62^ impeding the accumulation of Treg cells in VAT.^63^ Here, we demonstrate that, under physiological conditions, the Treg cell composition in AT mirrors that of other anatomical sites: pTreg cells represent 20-30% of the VAT-Treg compartment, with tTreg cells comprising 70-80%.

Although the pivotal role of VAT pTreg cells identified here was not anticipated based on prior research, their significance became evident early on in our investigations with Foxp3^RFP/GFP^ mice. When tTreg cells were selectively lost due to HFD feeding, pTreg cells failed to colonize the vacated niche (Figures 1H, 1I, and S2B), highlighting the stringent constraints imposed by the TCR-dependent niche. In response to the constitutive absence of tTreg cells, compensatory mechanisms enabled pTreg cells in ΔtTreg mice to occupy the VAT Treg cell niche, maintaining their functional specialization. This was further corroborated by a significant upregulation of genes previously identified as part of the ‘VAT- Treg’ signature in transcriptomic studies of total VAT Treg cells. Our finding that genetic pTreg ablation induces distinct ‘transcriptional holes’ in this ‘VAT-Treg’ signature in ΔpTreg mice underscores the non-redundant role of pTreg cells in VAT homeostasis. Moreover, it provides a mechanistic explanation for the spontaneous onset of pronounced obesity symptoms in ΔpTreg mice, even under normocaloric conditions. Consistent with this, HFD feeding exacerbated obesity in ΔpTreg mice, manifesting a severe phenotype spanning the full spectrum of T2D, including marked hepatic steatosis - an outcome rarely seen within 20 weeks of HFD feeding.^64,65^

The spontaneous onset of obesity symptoms following genetic pTreg cell ablation, as demonstrated here, mirrors observations in several other mouse mutants with disruptions in genes or pathways regulating energy balance and metabolism, such as leptin-deficient mice,^66,67^ pro- opiomelanocortin-deficient mice,^68^ and insulin receptor knockout mice.^69^ The only known protein with pleiotropic effects on VAT homeostasis in both non-immune (*e.g.*, adipocytes^70^) and immune cells^11,13,71–73^ is PPAR-γ. Genetic ablation of PPAR-γ in all cell types results in adipose tissue deficiency and a complex metabolic phenotype, including the development of T2D.^74^ Foxp3-dependent PPAR-γ ablation impedes the accumulation of Treg cells selectively in VAT, without affecting other tissues,^11,13^ with residual VAT PPAR-γ^−/−^ Treg cells exhibiting reduced Foxp3 expression and dysregulated ‘VAT Treg’ signature genes.^11,14^ Comparing Foxp3.PPAR-γ^−/−^ mice with the subset-specific loss-of-function models presented here shows that most phenotypic traits are similar to those observed in ΔpTreg mice. Both display heightened VAT inflammation^11,71^ and impaired glucose tolerance.^71^ However, the distinct roles of pTreg and tTreg cells revealed in our work likely account for phenotypic differences between these models. In particular, the rather moderate increase in VAT inflammation and attenuated body weight observed in Foxp3.PPAR-γ^−/−^ mice^71^ can be attributed to the absence of obesogenic tTreg cell transdifferentiation, a key driver of the exaggerated VAT inflammation and obesity seen in ΔpTreg mice - a phenomenon also abrogated in ΔtTreg mice. Furthermore, the ‘hyperplastic’ adipocyte morphology in Foxp3.PPAR-γ^−/−^ mice (*i.e.*, more, but smaller cells),^13^ juxtaposed with the shift to a ‘hypertrophic’ morphology in ΔtTreg mice (*i.e.*, fewer, but larger cells), indicates that the Treg cell-mediated regulation of adipocyte development^13^ is predominantly mediated by pTreg cells. Our findings suggest that the observed accumulation of adipogenic MSCs in the VAT of ΔtTreg mice is driven by an expanded VAT pTreg compartment expressing OSM. The ‘hypertrophic’ adipocyte morphology in these mice likely reflects a compensatory response to the restricted adipocyte differentiation mediated by pTreg cells. In line with our observation of improved glucose tolerance and insulin sensitivity in ΔtTreg mice fed a normocaloric diet, larger adipocyte size has been linked to improved metabolic performance.^75–78^ Taken together, these results underscore the indispensable role of VAT pTreg cells with a PPAR-γ^+^ST-2^+^ tissue phenotype in regulating VAT and metabolic homeostasis. Regarding their origin, our findings suggest that VAT pTreg cells are recruited from the splenic compartment of ST-2^+^ tissue-type Treg precursors, rather than being locally induced within the VAT itself. Future research will be critical to define the precise temporal and spatial dynamics of the recruitment and induction of tissue-type pTreg precursors, as well as to identify the antigens involved in this process.

The loss of VAT Treg cells, a hallmark of obesity, has previously been linked to adipocyte-related pro-apoptotic factors within a lipid-rich microenvironment. However, these studies did not explore whether pTreg and tTreg cells are differently affected. Our findings reveal differential expression of pro-survival factors (*e.g.*, *Pim1*, *Tnfaip3*, *Bcl2a1b*, *Bcl2*, *Mcl1*, and *Bcl2l1*) and metabolic fitness mediators (*e.g.*, *Hmgcr*, *Dgat1*, *Plin2*, and *Lmna*), with significant downregulation in the ‘Treg’ cluster of ΔpTreg mice. These results suggest that both apoptotic cell death and phenotypic destabilization likely contribute to the loss of VAT-Treg cells through non-exclusive mechanisms, which may share common initiating factors. Notably, dysregulated cytokine signaling pathways crucial for VAT Treg cell maintenance - such as selective loss of IL-2R and ST-2 expression, coupled with heightened pro-inflammatory cytokine pathways (TNF-α and IL-6) - appear central to these processes in VAT tTreg cells.

Given their clinical relevance for Treg cell-based therapies, the mechanisms governing the stability of mature Foxp3^+^ Treg cells have attracted significant attention in numerous studies. It is well-established that inflammatory and metabolic stress can trigger a cascade of events leading to the loss of Foxp3 expression and the disruption of Treg cell identity. This process includes the activation of inflammatory cytokine pathways, such as TNF-α and IL-6, which suppress Foxp3 expression,^79–81^ epigenetic alterations within the Foxp3 gene locus,^82–84^ miRNA-mediated regulation of Foxp3 mRNA,^85,86^ and increased cellular plasticity, including transdifferentiation into pathogenic Teff cell subsets.^81,87^ However, demonstrating the physiological significance of these processes has proven more challenging.

The loss of Treg cell identity *in vivo* and their transdifferentiation into pathogenic effector cells has been proposed to occur in a highly inflammatory environment,^81,82,87^ but this idea remains controversial.^55,56^ Notably, conclusions regarding Foxp3 loss *in vivo* were largely based on lineage tracing in experimental autoimmune models,^81,82,87^ an approach also used in our study, but contested by findings suggesting that Treg cells can remain remarkably stable in both steady-state and inflammatory conditions.^56^ Some studies even argue that the accumulation of Foxp3^−^YFP^+^ cells in peripheral tissues may be attributed to transient Foxp3 expression during thymic development rather than Foxp3 downregulation *in situ.*^55^ Here, we rule out this possibility by showing that VAT-derived Foxp3^−^YFP^+^ cells rapidly reacquire Foxp3 expression upon *in vitro* restimulation, a distinguishing feature that differentiates true ex-Treg cells from those that have already downregulated Foxp3 expression during thymic development. Additionally, while Treg plasticity after adoptive transfer into a lymphopenic environment has been shown to be limited to a subset of the Treg population - primarily enriched for naïve, Nrp-1− Treg cells, including pTreg cells and recent thymic emigrants^88^ - our flow cytometric and transcriptomic analyses suggest that this inherently unstable Treg subset is unlikely to significantly contribute to the loss of tTreg cell identity observed in obese VAT. Furthermore, we provide several lines of independent evidence supporting the origin of these cells from local tTreg dedifferentiation. This includes the identification of intermediate transdifferentiation stages (*e.g.*, *Foxp3* mRNA^low^ Teff cells), selective enrichment of Foxp3^−^YFP^+^ cells in VAT (but not in SAT or lymphoid tissues), and TCR clonotype trajectory analyses, which show the accumulation of unique tTreg clonotypes specifically in the pro-inflammatory Th1 compartment, but not in Th2.

The pro-inflammatory IFN-γ^+^ phenotype of Foxp3^−^YFP^+^ cells, coupled with their marked correlation with the severity of obesity, strongly suggests that ex-tTreg cells exacerbate obesogenic inflammation. This hypothesis is further supported by genetic tTreg cell ablation, which prevented the accumulation of ex-tTreg clonotypes and mitigated obesity symptoms. While our studies primarily focused on the selective loss of VAT tTreg cells, the transdifferentiation of these cells into obesogenic Teff cells, along with the compensatory actions of pTreg cells, may have obscured key biological functions of tTreg cells. According to the prevailing view, tTreg cells are predominantly selected by self-antigens during intrathymic development and are functionally specialized to maintain immune homeostasis and prevent autoimmune responses.^89,90^ This concept is further reinforced by the Foxp3^RFP/GFP^ model, in which selective tTreg cell deficiency on the genetically predisposed NOD background leads to an especially severe form of type 1 diabetes.^29^ Although adipose tissues are not typically viewed as primary targets of autoimmune responses, installing TCRs specific to VAT-restricted self-antigens into the tTreg cell pool could offer a mechanism for directing these cells to the VAT, where they might carry out specialized functions within the local microenvironment.^15,28,91^ At this stage, we can only hypothesize about the biological significance of tTreg cell transdifferentiation within the obese VAT. The ability of ex-tTreg cells to reacquire Foxp3 expression may serve as an adaptive immune strategy to restore tissue homeostasis, neutralize danger signals (such as toxic lipids and metabolites), and promote tissue repair through controlled inflammation.^92–95^ However, for this process to be effective, it requires a precisely timed response to avoid collateral damage, particularly in conditions of chronic immunological and metabolic stress, such as prolonged high-calorie intake that compromise tTreg cell stability.

Our studies have centered on elucidating the roles of pTreg and tTreg cells in maintaining VAT homeostasis under both steady-state and obesogenic conditions. Future studies are crucial to uncover the precise mechanisms by which VAT Treg cells interact with EOs,^33^ whose population size declines alongside tTreg cells, and well as other immune populations regulating VAT homeostasis, such as group 2 innate lymphoid cells,^96^ and invariant natural killer T cells.^97–99^ Given the increasing global burden of metabolic disorders, these insights hold promising potential for the rational development of novel therapeutic strategies.

## Limitations of Study

Our experiments reveal distinct roles of the two VAT Treg cell developmental subsets in tissue homeostasis and obesogenic inflammation, demonstrating that these processes are driven by divergent TCR specificities. However, the precise antigens involved remain unidentified. Characterizing these antigens is pivotal for bridging critical gaps in our understanding of the mechanisms underlying this functional specialization. An unresolved question persists as to whether the TCR specificities of pTreg- and tTreg-derived VAT Treg cells modulate VAT inflammation and metabolic pathways in accordance with the distinct roles identified in this study. Addressing this will necessitate complex experiments, including single-cell TCR sequencing across various anatomical sites (thymus, colon, spleen, lymph nodes, adipose tissues), identification and isolation of relevant TCRs, and the generation of TCR transgenic mouse lines to explore fate decisions and functional outcomes *in vivo*. Such studies will also further clarify the relationship between splenic tissue Treg precursors and VAT Treg developmental subsets. Although demanding, these efforts are invaluable, offering the potential to locally modulate VAT Treg cell numbers and activity in an antigen- and subset-specific manner.

## SUPPLEMENTAL INFORMATION (11 figures and 2 tables)

### Supplementary Figures

Figure S1 (related to Figure 1). Flow cytometry of immune cells in Foxp3RFP/GFP mice

Figure S2 (related to Figure 1). Flow cytometry of T cells in Foxp3RFP/GFP mice

Figure S3 (related to Figure 2). AT weight and multicolor flow cytometry of innate immune cells in ΔtTreg mice after 20 weeks of HFD feeding

Figure S4 (related to Figure 3). HFD exacerbates obesity in ΔpTreg mice

Figure S5 (related to Figure 4). CD4+ T cell responses in VAT of ΔtTreg and ΔpTreg mice

Figure S6 (related to Figure 4). Differential gene expression of key markers in VAT CD4+ T cell clusters Figure S7 (related to Figure 5). Stromal adipocyte precursor cells in ΔtTreg and ΔpTreg mice

Figure S8 (related to Figure 6). VAT Treg cell surface markers in HFD-fed ΔtTreg and ΔpTreg mice

Figure S9 (related to Figure 6F). Unstable VAT Treg cells in the ’Tmem’ cluster of ΔpTreg mice Figure S10 (related to Figure 7). Characterization of VAT ex-tTreg cells

Figure S11 (related to Figure 4). Isolation of VAT CD4+ T cells for scRNA/TCR-seq

## Supplementary Tables

Table S1. Top 50 differentially expressed genes among VAT CD4^+^ T cell clusters (see Figures 4D and S6) Table S2. Top 50 differentially expressed genes among ’Tmem’ sub-clusters (see Figures 6F and S9)

## ACKNOWLEDGMENTS

The authors would like to thank Lena Biedermann (CRTD), as well as and Bettina Gercken and Christine Mund (Institute for Clinical Chemistry and Laboratory Medicine) for technical assistance; and Katja Bernhardt and Anne Gompf (CMCB Flow Cytometry Core Facility) for expert help in flow cytometry and cell sorting.

## AUTHOR CONTRIBUTIONS

AY: Writing – original draft, Conceptualization, Data curation, Formal analysis, Investigation, Methodology, Visualization. AE, DMZ, SSH, MB, BT, EM, EK, AP, OK: Writing – review & editing, Formal analysis, Investigation, Methodology, Visualization. AD: Writing – review & editing, Conceptualization, Formal analysis, Resources. VIA: Writing – review & editing, Conceptualization, Formal analysis. AC: Writing – review & editing, Conceptualization, Funding acquisition, Methodology, Project administration, Resources, Supervision. MD: Writing – original draft, Conceptualization, Data curation, Formal analysis. SS: Writing – original draft, Conceptualization, Formal analysis, Resources. KK: Writing – original draft, Conceptualization, Data curation, Formal analysis, Funding acquisition, Investigation, Methodology, Project administration, Resources, Supervision, Validation.

## METHODS

### Mouse models

C57BL/6 (B6) WT mice were obtained commercially (C57BL/6NRj; Janvier Labs). Foxp3^RFP/GFP^,^25^ R26Y x Foxp3^RFP/GFP^,^31^ R26^DTA^ x Foxp3^RFP/GFP^ (ΔtTreg^29^), and Foxp3-STOP x Foxp3^RFP/GFP^ (ΔpTreg^29^) mice were generated on the B6 background. All mouse lines were bred and housed under specific-pathogen-free (SPF) conditions at the Animal Facility of the CRTD. Mice were fed a VegND (9 kcal% fat from soybean oil, 24 kcal% protein, and 67 kcal% carbohydrates; Ssniff, Germany). To induce obesity, cohorts of 6- week-old males with the indicated genetic modifications were fed a HFD (60 kcal% fat, 5.55% soybean oil, 54.35% lard; 20 kcal% protein, 20 kcal% carbohydrates; Research Diets, USA). Control mice were fed an ND (10 kcal% fat, 5.55% soybean oil, 4.44% lard; 20 kcal% protein, 70 kcal% carbohydrates; Research Diets, USA). Body weight was measured before and weekly after the initiation of feeding experiments. Animal experiments were conducted in accordance with approvals from the Landesdirektion Dresden (25-5131/502/5, TVA 5/2020; 25-5131/522/43, TVV41/2021; 25- 5131/562/5, TVV 3/2023; 25-5131/564/21, TV vG 20_2023).

### Histopathology

Histopathological analyses of the small intestine and mesenteric AT were performed as previously described.^29^ For histological examination of SAT, VAT, and liver, 26- to 27-week-old mice were used, either following 20 weeks of HFD feeding or VegND. After euthanizing the mice with CO2 inhalation, the left ventricle and portal vein were perfused with 50 ml of 1x PBS (ThermoFisher, Life Technologies). Organs were collected, briefly washed in PBS, and fixed in 4% paraformaldehyde (PFA) solution (Sigma-Aldrich) before paraffin embedding. 4 µm sections were cut, and four sections from each sample were collected at 50 μm intervals and stained with hematoxylin and eosin for morphological analysis. Inflammation in SAT and VAT was scored blindly as follows: Score 0 (no immune cell infiltration), Score 1 [minor number of crown-like structures (CLS)], Score 2 (moderate number of CLS), Score 3 (extensive infiltration beyond peri-adipocyte area). Median adipocyte size and number were measured using the ImageJ AdipoQ plugin^100^ in five 1500 x 800 µm² areas and six 750 x 400 µm² areas for HFD- and VegND-fed mice, respectively. Liver steatosis was graded blindly based on the percentage of fat within hepatocytes: Score 1 (<60%), Score 2 (60-84%), Score 3 (>85%).

### Insulin and glucose tolerance test

Metabolic tests were conducted in weeks 18 and 19 of HFD feeding or in age-matched VegND-fed mice, as specified. For the glucose tolerance test (GTT), mice underwent a 6-hour fast before receiving an intraperitoneal (*i.p*.) injection of glucose (1 g/kg body weight). For the insulin tolerance test (ITT), insulin (1 U/kg for VegND-fed and 1.5 U/kg for HFD-fed mice; Huminsulin, Lilly) was administered *i.p.* after a 6-hour fast. Blood glucose levels were measured at baseline and at different timepoints post-injection (15, 30, 60, and 90 minutes). The area under the curve (AUC) was calculated for each mouse.

### Flow cytometry

#### Preparation of Single-Cell Suspensions from lymphoid tissues

Single-cell suspensions were prepared in Hank’s buffer [1× HBSS, 5% (v/v) FCS, 10 mM HEPES; all ThermoFisher, Life Technologies]. Spleen was passed through 70 μm cell strainers (BD Biosciences). Single-cell suspensions from spleen were then treated with erythrocyte lysis buffer (EL; Qiagen) to remove red blood cells.

#### Isolation of SVF from AT

The left ventricle and portal vein were perfused with 50 ml of 1× PBS (ThermoFisher, Life Technologies) as previously described.^7^ After removing superficial inguinal lymph nodes from the SAT and detaching VAT from the testes and remaining reproductive tissues, the tissues were kept in 1× PBS supplemented with 1 mM CaCl₂ and 0.5% (w/v) bovine serum albumin (BSA) (Sigma Aldrich/Merck) on ice. The tissues were minced into small pieces and digested with a 2 mg/ml collagenase solution (Collagenase from *Clostridium histolyticum* Type I, Thermo Fisher Scientific) dissolved in high-glucose DMEM (Thermo Fisher, Life Technologies) containing 0.5% (w/v) BSA at 37°C for 60 minutes using a thermomixer (Eppendorf).^101^ After digestion, the mixture was filtered through a 100 μm cell strainer (Corning) and centrifuged. Floating adipocytes were separated from the SVF, which was treated with erythrocyte lysis buffer (EL buffer; Qiagen), dissolved in Hank’s buffer, and filtered through a 30 µm cell strainer (Sysmex Partec). Immune cells isolated from SAT and VAT were stored separately for downstream applications.

#### Flow cytometry and cell sorting

Monoclonal antibodies (mAbs) to CCR2 (475301), CD3ε (145-2C11), CD4 (RM4-5), CD8a (53-6.7), CD11b (M1/70), CD11c (HL3), CD14 (Sa2-8), CD16/32 (93), CD19 (1D3), CD25 (PC61), CD31 (390), CD44 (IM 7), CD45.1 (A20), CD45.2 (104), CD62L (MEL-14), CD64 (X54-5/7.1), CD103 (M290), CD127 (A7R34), CD206 (MR5D3), F4/80 (BM8), GITR (DTA-1), ICOS (7E.17G9), KLRG1 (2F1), LY-6G (1A8), OX-40 (OX-86), PD-1 (29F.1A12), Pdgfrα (APA5), Sca-1 (D7), Siglec-F (E50-2440), ST- 2 (U29-93), Thy1.2 (30-H12), Foxp3 (FJK-16s), GATA-3 (L50-823), IFN-γ (XMG1.2), IL-10 (JES5-16E3), IL-17a (eBio17B7), IL-2 (JES6-5H4), IL-4 (11B11), IL-5 (TRFK5), TNF (MP6-XT22), Fc receptor-blocking mAb against CD16/32 (93), and fluorochrome-conjugated streptavidin (BUV395, eFlour450, APC and PE- Cy7) were purchased from BD, eBioscience, or BioLegend. mAb to PPAR-γ (4E12F10) was purchased from Proteintech. Pre-gating included CD45 gating and FSC/SSC-based doublet-exclusion for all immune cells, followed by non-T cell-exclusion for T cell analyses using an ‘exclusion mix’ (EM) of mAbs directed against CD14, CD11b, CD11c, and CD19. Intracellular cytokine and transcription factor expression was analyzed using fluorochrome-conjugated mAbs in combination with the BD Cytofix/Cytoperm kit (BD) and the Foxp3 Staining Buffer Set (eBioscience), following the manufacturer’s protocols. Viable cell numbers were determined using propidium iodide and a MACSQuant (Miltenyi Biotec). Prior to CD4^+^ T cell isolation from VAT via FACS, samples were depleted of lineage marker-positive cells (CD8a, CD11b, CD11c, CD14, and CD19) using biotin-conjugated mAbs, along with streptavidin-conjugated microbeads and the autoMACS Pro magnetic separation system (Miltenyi Biotec). Samples were stained with DRAQ7 (BioStatus) for dead cell exclusion, filtered through 40 µm cell strainers, and analyzed on an LSR Fortessa or sorted with a FACS Aria II or III (all BD). Data were analyzed using FlowJo software (Version 10.8.1, Tree Star Inc.).

#### T cell culture

Unless otherwise stated, T cells were cultured in 96-well round-bottom plates (Greiner) at 37°C and 5% CO₂ in 200 μl RPMI complete medium [RPMI 1680 medium supplemented with 1 mM sodium pyruvate, 1 mM HEPES, 2 mM Glutamax, 100 U/ml penicillin, 100 µg/ml streptomycin, 100 µg/ml gentamycin, 0.1 mM non-essential amino acids, 0.55 mM β-mercaptoethanol, and 10% FCS (v/v); all ThermoFisher, Life Technologies]. For intracellular cytokine analysis by flow cytometry, single-cell suspensions were stimulated for 4 hours in RPMI complete medium with 50 ng/ml phorbol 12- myristate 13-acetate (PMA) and 200 ng/ml ionomycin (Iono), in the presence of 20 μg/ml Brefeldin A (all Merck, Sigma-Aldrich). T cells, including CD4^+^CD25^−^YFP^−^RFP^−^ Tcon, CD4^+^YFP^+^RFP^+^ tTreg, and CD4^+^YFP^+^RFP^−^ ex-tTreg cells, were sorted from CD4-bead-enriched single-cell suspensions of VAT from R26Y x Foxp3^RFP/GFP^ mice fed HFD for 20 weeks. Prior to culture initiation, 96-well flat-bottom plates (Greiner) were coated overnight at 4°C with 10 µg/ml anti-CD3ε mAb (145-2C11, BD) diluted in 1× PBS. The sorted cells were cultured in the anti-CD3ε-coated plates with RPMI complete medium containing soluble anti-CD28 mAb (2 µg/ml, 37.51, eBioscience) and recombinant human IL-2 (1000 U/ml, Teceleukin, Roche). On day 3, the expression of CD25, YFP, and RFP was assessed by flow cytometry.

#### Protein extraction and immunoblotting

ATs were shock-frozen using dry ice and homogenized with a Pequlab Precellys 24 Homogenizer in lysis buffer [1% Triton X-100, 0.5% sodium deoxycholate, 0.1% SDS, 50 mM Tris-HCl (pH 7.5), 150 mM NaCl; all Sigma Aldrich/Merck], supplemented with a protease inhibitor and phosphatase inhibitor cocktail (Roche). The lysates were incubated in an ultrasonic bath. Protein concentrations were measured using the BCA Protein Assay Kit (Thermo Scientific). For immunoblotting, 10 µg of total protein from each sample were loaded. The blots were probed with primary antibodies against phospho-ERK1/2 and total-ERK1/2 (Cell Signaling/New England Biolabs), followed by anti-rabbit HRP-conjugated secondary antibodies. Signals were developed using SuperSignal West Pico Chemiluminescent Substrate (Thermo Scientific) and detected with a LAS-3000 luminescent image analyzer (Fujifilm).

#### Single-cell transcriptome and TCR profiling

##### Sample preparation

For single-cell RNA and TCR sequencing (scRNA/TCR-seq), B6.Foxp3^RFP/GFP^ and B6 mice with selective pTreg and tTreg cell deficiency were fed a HFD for 20 weeks. Single-cell suspensions of VAT from ≥ 4 mice were pooled for each experimental condition. Following magnetic bead-based lineage depletion, 1.5 × 10⁴ viable CD45^+^CD4^+^ cells were FACS-purified using a 70 µm nozzle and the 4-way purity sort mode. The sorted cells were collected into 5 µl PBS in 1.5 ml Eppendorf tubes, which were coated overnight with 1% (w/v) BSA (Sigma Aldrich/Merck).

##### scRNA/TCR-seq library preparation and sequencing

Samples were processed at the Deep Sequencing Facility of the DRESDEN-concept Genome Center at the CRTD. Briefly, sorted CD4^+^ T cells were immediately loaded onto the 10x Chromium chip. The Single Cell 5’ reagent kit [Chromium Next GEM Single Cell V(D)J Reagent Kit v1.1] was used for reverse transcription, cDNA amplification, and construction of gene expression libraries, following the protocol provided by 10x Genomics. Libraries were sequenced on a NovaSeq 6000 S4 v1.5 flowcell in XP mode with 100 bp paired-end reads, achieving an average sequencing depth of 50K reads per cell for gene expression and 5K reads for each TCR- and feature barcode.

##### Analysis of scRNA-seq data

Raw sequencing data were processed using the Cell Ranger software (v2.1.0) from 10x Genomics. The mouse genome (mm10) and gene annotations (v87) were downloaded from Ensembl, and the annotation was filtered with the ’mkgtf’ command of Cell Ranger (--attribute = gene_biotype:protein_coding –attribute = gene_biotype:lincRNA –attribute = gene_biotype:antisense). The genome sequence and filtered gene annotations were used to build the appropriate Cell Ranger reference. Downstream analysis was performed using the Seurat package^102^ in R (version 3.6.2). Cells were excluded if they expressed fewer than 500 or more than 2500 unique genes, or if mitochondrial transcripts exceeded 4%. Genes not detected in any cell were excluded from subsequent analysis. After quality control, 17,120 features across 24,653 high-quality cells (Foxp3^RFP/GFP^ mice: 9091 cells; ΔtTreg mice: 7934 cells; and ΔpTreg mice: 7628 cells) were obtained and used for downstream transcriptional analysis.

The data were log-normalized using *NormalizeData*, and highly variable genes were identified with *FindVariableGenes* using default parameters. Cell cycle scores were assigned to each cell using *CellCycleScoring* and regressed out during data scaling with the *ScaleData* function, including the *vars.to.regress* option. Principal component (PC) analysis was performed on the scaled data, and the 14 most contributing PCs were used for downstream analyses of all cells. For sub-clustering of ’Tmem’ cluster, the 10 most contributing PCs were used. For dimensionality reduction, UMAP was applied using *RunUMAP* and visualized with *DimPlot*. Differentially expressed genes for each cluster/sub- cluster were identified using *FindAllMarkers* and used to define cluster/sub-cluster identities. Main cluster identities were determined based on differential gene expression of canonical markers. For the ’Treg’ and ’Tmem’ clusters, 2526 (Foxp3^RFP/GFP^ mice: 1116 cells; ΔtTreg mice: 900 cells; ΔpTreg mice: 510 cells) and 4386 cells (Foxp3^RFP/GFP^ mice: 1273 cells; ΔtTreg mice: 1216 cells; ΔpTreg mice: 1897 cells) were obtained, respectively, and used for downstream analyses.

The expression of *Cd8a*, *Cd49a*, and *Cd19* was not found in any sequenced cells showing the absence of CD8^+^ T, NK, and B cells, and the purity of sorted cells, which were confirmed by post-sort analyses (Figure S11A). The CD4^+^ T cell identity of sequenced cells was further validated by visualization of widespread expression of *Ptprc* (CD45), *CD3e* (CD3ε), and *Cd4* (CD4) (Figure S11B), and by the virtual absence of expression of canonical markers for other immune cells [Monocytes, Mφs, granulocytes, and DCs: *Itgam* (CD11b), *Itgax* (CD11c); B cells: *Cd79a* (CD79a); NK cells and ILCs: *Klrb1c* (NK1.1); and γδ T cells: *Trdc* (TRDC)] and nonimmune cells [MSCs: *Pdgfra* (PDGFRα), endothelial cells: *Cd34* (CD34), pericytes: *Mcam* (CD146), and adipocytes: *Adipoq* (Adiponectin)]^103,104^ (Figure S11C).

##### Analysis of scTCR-seq data

TCR sequences were extracted from 10x Genomics data using Cell Ranger (“cellranger vdj”, 10x Genomics). 10x Genomics TCR data were further analyzed using the scRepertoire package^105^ and mapped to the expression data with the “combineTCR/BCR” function. Clonotypes were matched by the VDJC segments of both *Tcra* and *Tcrb* with identical *Cdr3* sequences at the nucleotide level, if not stated otherwise. The figures for shared TCR clonotypes and the clonal abundance pie charts were generated using ggplot2.

#### Statistical analysis

Statistical significance was assessed using Prism 8 software (Version 8.4.3, GraphPad Software Inc., CA, USA). As indicated, the Student’s *t*-*test* (unpaired, two-tailed), two-way ANOVA (multiple comparisons with Sidak correction), non-linear regression with comparison of fits, and Pearson correlation analysis were used to assess statistical significance. Differences were considered as significant when *p ≤ 0.05, **p ≤ 0.01, ***p ≤ 0.001, ****p ≤ 0.0001.

