## Supplemental Information for "Functional Dichotomy of Developmental Foxp3^+^ Treg Cell Subsets in the Visceral Adipose Tissue of Lean and Obese Mice"

### Figure S1 (related to Figure 1). Flow cytometry of immune cells in Foxp3$^{RFP/GFP}$ mice

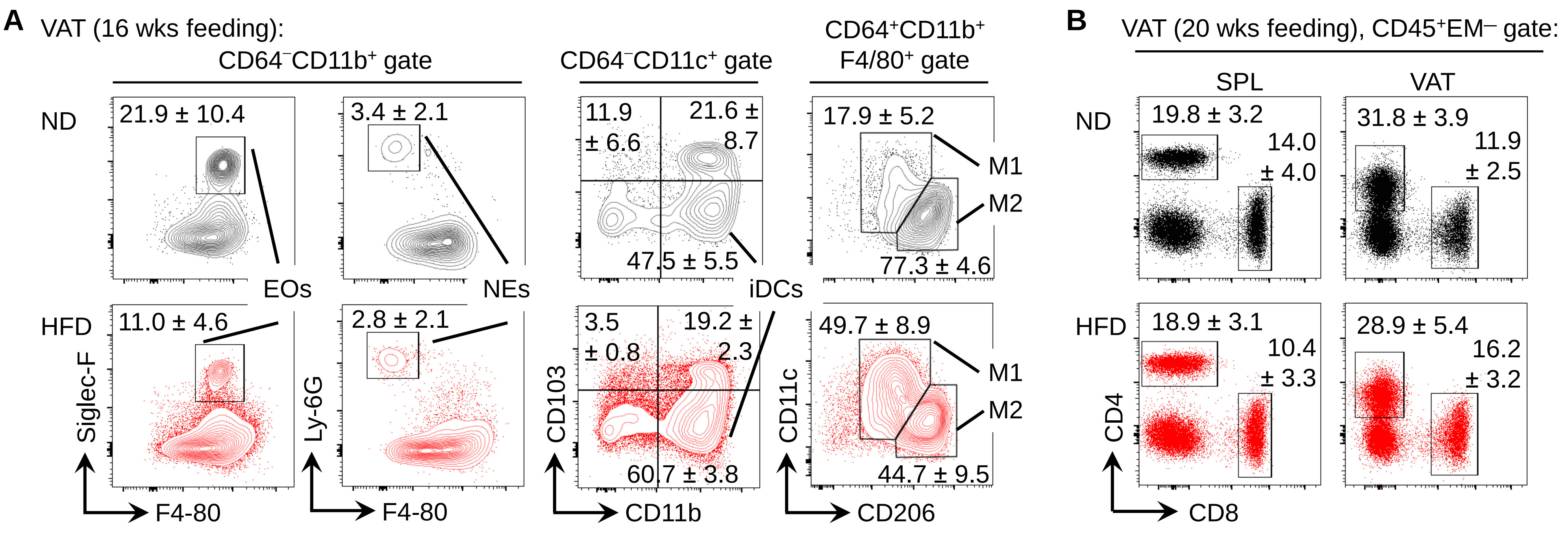


**Figure S1 (related to Figure 1). Flow cytometry of immune cells in Foxp3$^{RFP/GFP}$ mice**

(A) Contour plots of indicated innate immune cell subsets in VAT after 16 weeks of ND (top) or HFD (bottom). NEs, neutrophils; iDCs, inflammatory DCs; M1/M2, macrophage subsets.

(B) Dot plots of CD4$^{+}$ and CD8$^{+}$ T cells in spleen (SPL, left) and VAT (right) after 20 weeks of ND (top) or HFD (bottom).

Mean percentages ± SD. Representative experiment (n = 6) of three independent experiments (n = 3 - 6 mice/experiment).

**Figure S2 (related to Figure 1). Flow cytometry of T cells in Foxp3$^{RFP/GFP}$ mice**

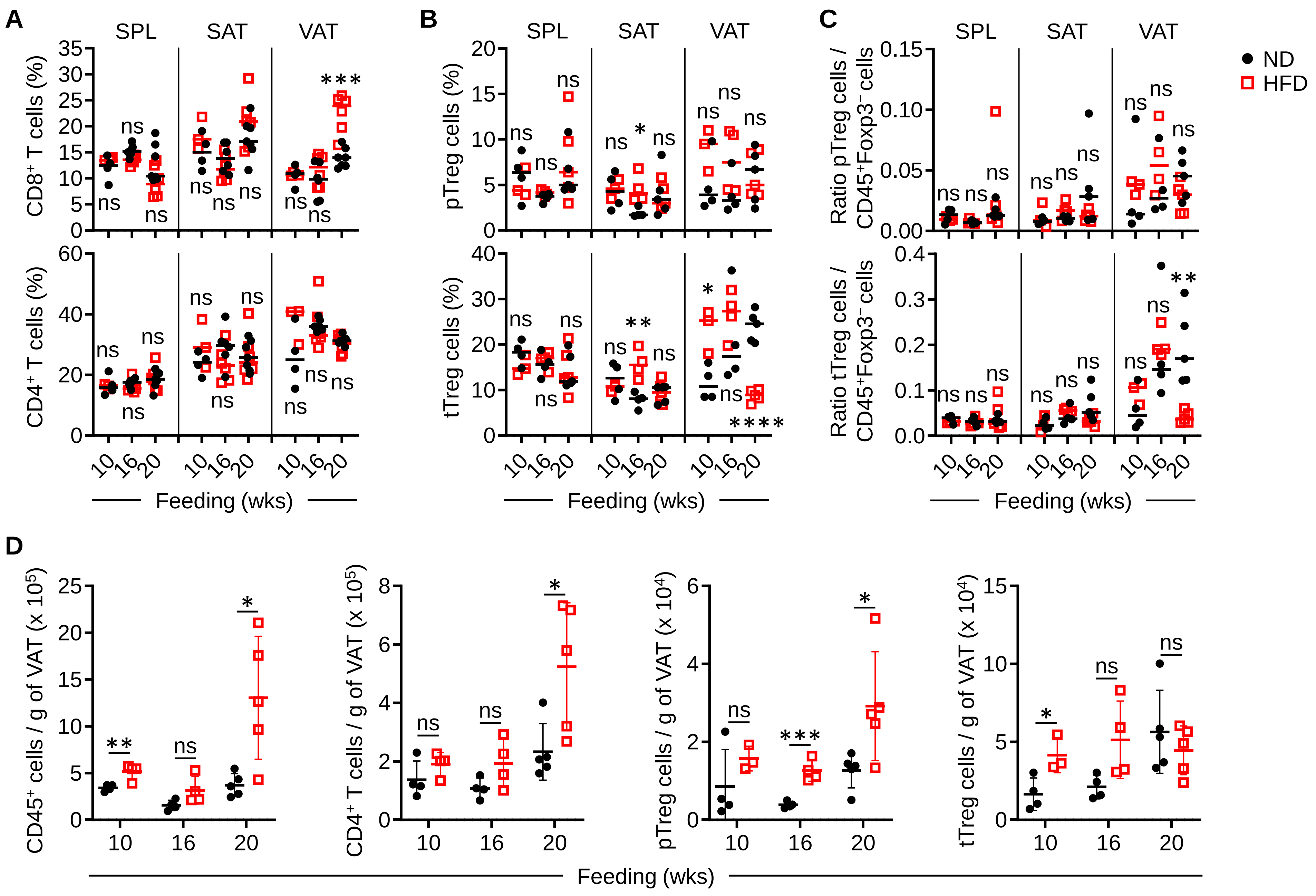


**Figure S2 (related to Figure 1). Flow cytometry of T cells in Foxp3$^{RFP/GFP}$ mice**

(A and B) Cumulative percentages of T cell subsets in spleen (SPL), SAT, and VAT after 10, 16, and 20 weeks of ND (top) or HFD (bottom).

(A) CD8$^+$ (top) and CD4$^+$ (bottom) T cells.

(B) RFP$^+$GFP$^-$ pTreg (top) and RFP$^+$GFP$^+$ tTreg (bottom) cells.

(C) Ratio of pTreg (top) and tTreg (bottom) cells to CD45$^+$Foxp3$^-$ cells in spleen (SPL), SAT, and VAT.

(D) Numbers of CD45$^+$, CD4$^+$, pTreg and tTreg cells normalized to VAT weight (g).

Symbols and lines indicate individual mice and mean values, respectively. Unpaired t-test: ns, not significant; *p ≤ 0.05, **p ≤ 0.01, ***p ≤ 0.001, ****p < 0.0001. Representative experiment (n = 3 – 6 per group) from three independent experiments (n = 3 – 6/experiment).

**Figure S3 (related to Figure 2). AT weight and multicolor flow cytometry of innate immune cells in ΔtTreg mice after 20 weeks of HFD feeding**

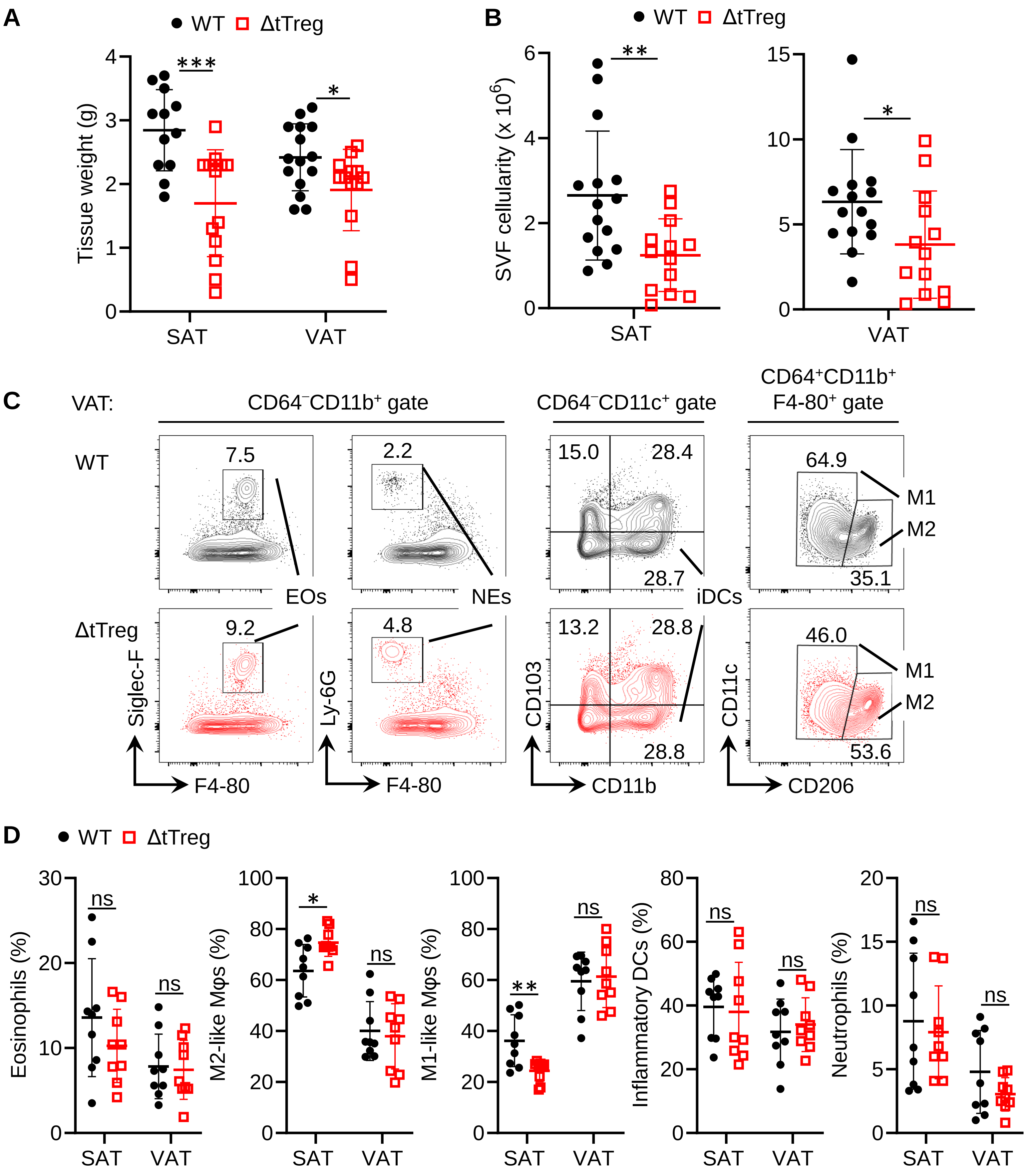


**Figure S3 (related to Figure 2). AT weight and multicolor flow cytometry of innate immune cells in ΔtTreg mice after 20 weeks of HFD feeding**

(A and B) Reduced (A) AT weight and (B) SVF cellularity in ΔtTreg mice (n = 13) compared to Foxp3$^{RFP/GFP}$ (WT, n = 15) mice.

(C and D) Attenuated HFD-induced shift to proinflammatory type 1 immunity in ΔtTreg mice.

(C) Contour plots show innate immune cell subsets in VAT of Foxp3$^{RFP/GFP}$ (WT, top) and ΔtTreg (bottom) mice. Percentages within the respective gate or quadrant.

(D) Cumulative percentages of innate immune cells in SAT and VAT (n = 9/group). NEs, neutrophils; iDCs, inflammatory DCs; M1/M2, macrophage subsets.

Graphs show individual mice and mean values ± SD (error bars). Unpaired t-test: ns, not significant; *p ≤ 0.05, **p ≤ 0.01, ***p ≤ 0.001 (A, B, and D). Data representative of three independent experiments.

**Figure S4 (related to Figure 3). HFD exacerbates obesity in ΔpTreg mice**

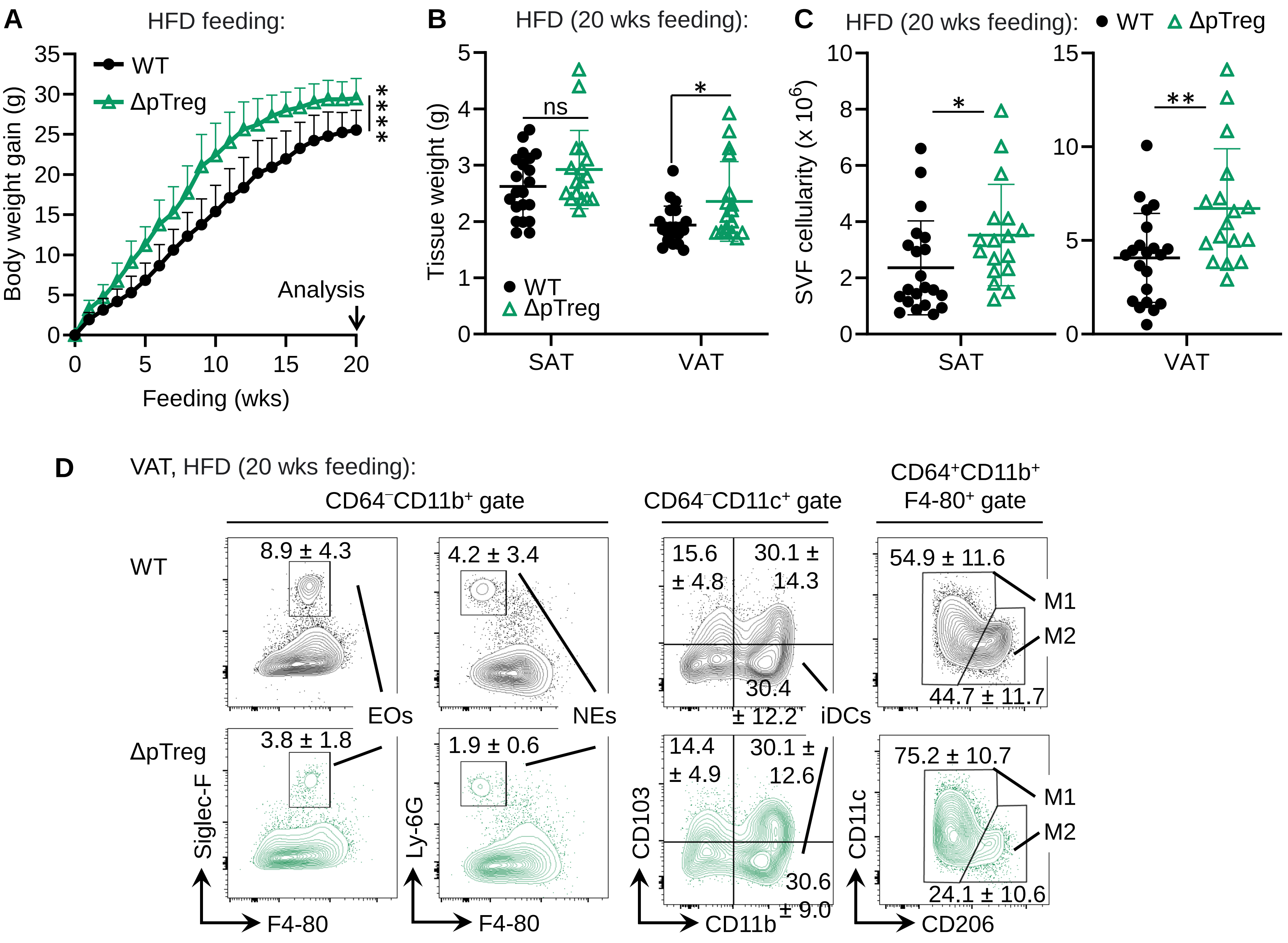


**Figure S4 (related to Figure 3). HFD exacerbates obesity in ΔpTreg mice**

(A-C) Increased body and AT weight in ΔpTreg mice after 20 weeks of HFD.

(A) Body weight gain in Foxp3$^{RFP/GFP}$ (WT, n = 21) and ΔpTreg (n = 17) mice. Mean ± SD. Two-way ANOVA with Sidak's correction; 20 weeks: ****p ≤ 0.0001.

(B) AT weight and (C) SVF cellularity. Graphs show individual mice (symbols) and mean ± SD (error bars). Unpaired t-test: ns, not significant; *p ≤ 0.05, **p ≤ 0.01.

(D) Enhanced proinflammatory type 1 shift in VAT of ΔpTreg mice. Contour plots show innate immune cell subsets in VAT of Foxp3$^{RFP/GFP}$ (WT; n = 6) and ΔpTreg (n = 7) mice. NEs, neutrophils; iDCs, inflammatory DCs; M1/M2, macrophage subsets. Numbers indicate mean percentages ± SD. Data from two experiments, representative of three independent experiments (n = 3 - 4 mice/experiment).

**Figure S5 (related to Figure 4). CD4⁺ T cell responses in VAT of ΔtTreg and ΔpTreg mice**

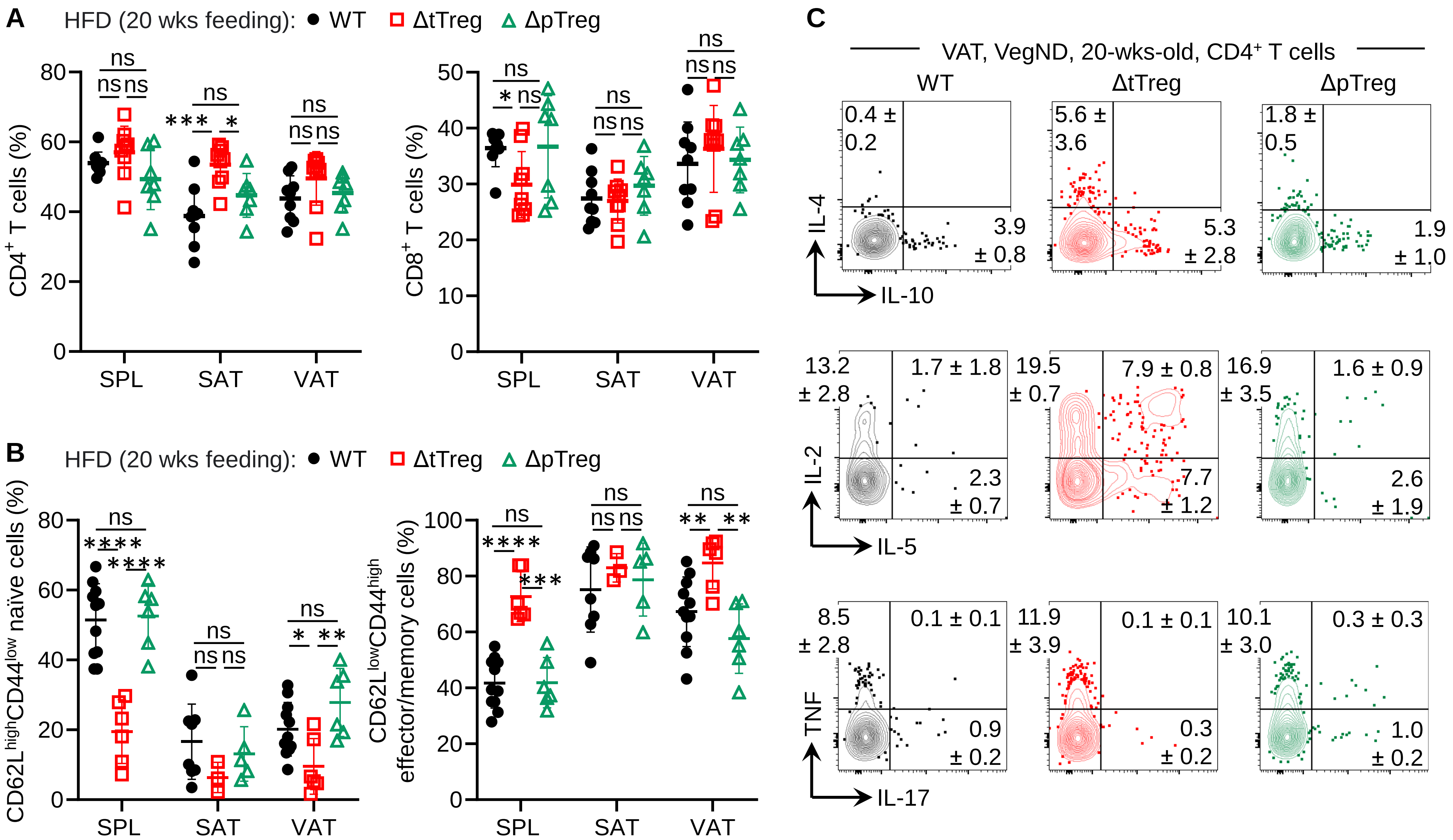


**Figure S5 (related to Figure 4). CD4⁺ T cell responses in VAT of ΔtTreg and ΔpTreg mice**

(A-B) Flow cytometry of VAT immune cells in $Foxp3^{RFP/GFP}$, ΔtTreg, and ΔpTreg mice fed HFD for 20 weeks (Figures 4A and B). Composite percentages of (A) $CD4^+$ (left) and $CD8^+$ (right) T cells, and (B) naïve ($CD62L^{high}CD44^{low}$, left) and effector/memory ($CD62L^{low}CD44^{high}$, right) $CD4^+Foxp3^-$ T cells. Graphs show individual mice (symbols) and mean ± SD (error bars). Unpaired t-test: ns, not significant; *p ≤ 0.05, **p ≤ 0.01, ***p ≤ 0.001, ****p ≤ 0.0001. Data from three independent experiments (6 - 11 mice/group).

(C) Representative flow cytometry of intracellular cytokine production in VAT $CD4^+$ T cells from VegND-fed $Foxp3^{RFP/GFP}$ (WT), ΔtTreg, and ΔpTreg mice (Figure 4F). Mean ± SD. Data from a single experiment (n = 3 - 5 mice/group).

### Figure S6 (related to Figure 4). Differential gene expression of key markers in VAT CD4+ T cell clusters

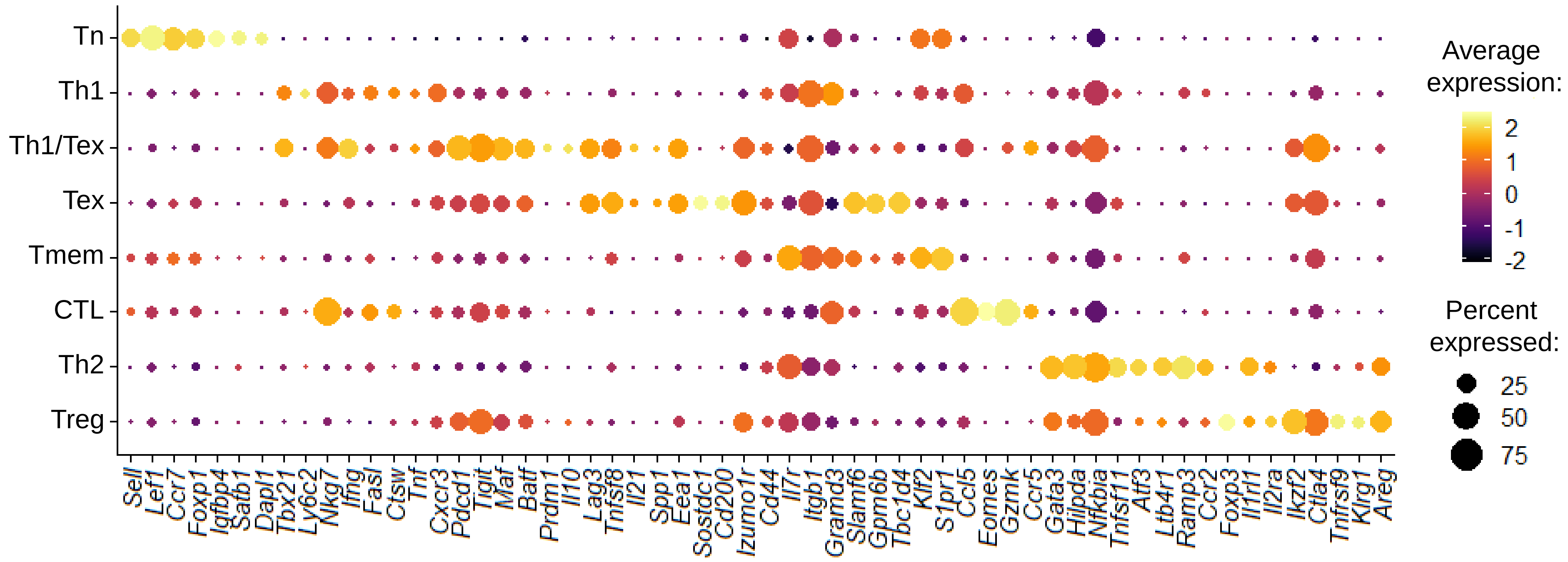


**Figure S6 (related to Figure 4). Differential gene expression of key markers in VAT CD4+ T cell clusters**
Bubble plot depicting the average and percent expression of key cluster-defining genes across eight distinct VAT CD4+ T cell clusters from HFD-fed Foxp3$^{RFP/GFP}$, ΔtTreg, and ΔpTreg mice (Figure 4D). Key markers were selected from the top 50 differentially expressed genes listed in Supplementary Table S1.

### Figure S7 (related to Figure 5). Stromal adipocyte precursor cells in ΔtTreg and ΔpTreg mice

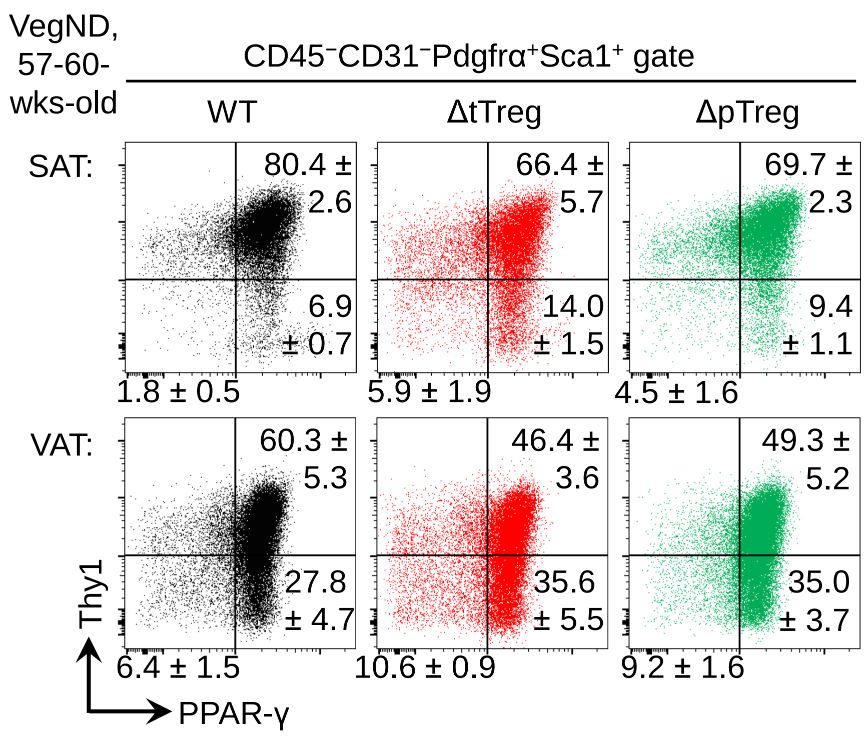


**Figure S7 (related to Figure 5). Stromal adipocyte precursor cells in ΔtTreg and ΔpTreg mice**

Flow cytometry of stromal adipocyte precursor cells in Foxp3$^{RFP/GFP}$ (WT, n = 4), ΔtTreg (n = 5), and ΔpTreg (n = 3) females fed VegND. Representative dot plots of Thy1 and PPAR-γ expression among gated CD45$^{-}$CD31$^{-}$Pdgfrα$^{+}$Sca1$^{+}$ MSCs in SAT (top) and VAT (bottom) of WT (left), ΔtTreg (middle), and ΔpTreg (right) mice. Numbers indicate percentages of cells ± SD within each gate. Composite percentages of adipogenic MSC subsets are shown in Figure 5H. Data are from a single experiment, representative of two independent experiments (3 - 5 mice/group).

**Figure S8 (related to Figure 6). VAT Treg cell surface markers in HFD-fed ΔtTreg and ΔpTreg mice**

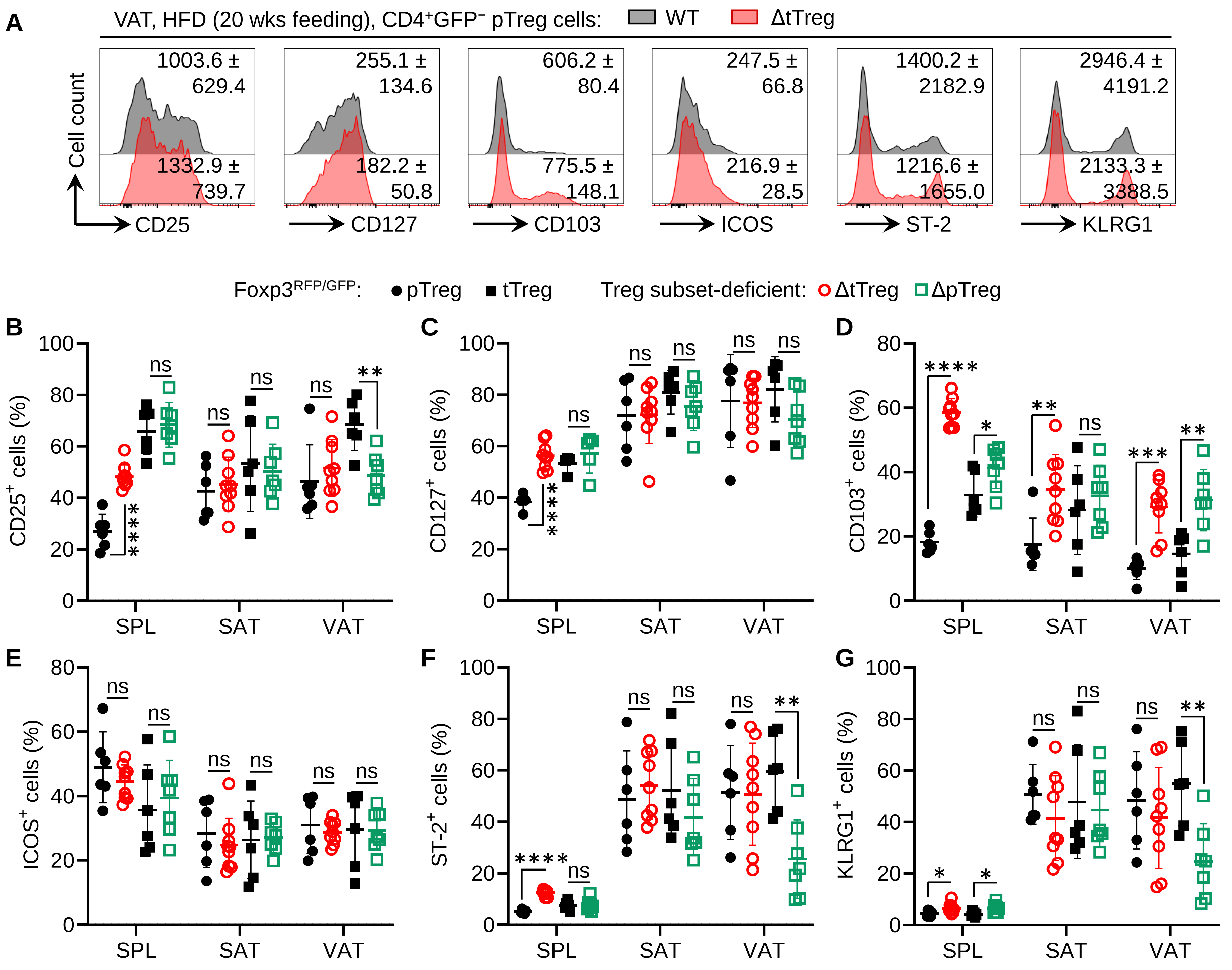


**Figure S8 (related to Figure 6). VAT Treg cell surface markers in HFD-fed ΔtTreg and ΔpTreg mice**

Flow cytometry of VAT Treg cells in $Foxp3^{RFP/GFP}$ (WT), ΔtTreg, and ΔpTreg mice after 20 weeks of HFD.

(A) Histograms of selected surface marker expression in VAT pTreg cells ($CD4^{+}RFP^{+}GFP^{-}$) from WT (grey) and ΔtTreg (red) mice. Mean MFI ± SD.

(B-G) Cumulative percentages of (B) CD25, (C) CD127, (D) CD103, (E) ICOS, (F) ST-2, and (G) KLRG1 expression among Treg cells in spleen (SPL), SAT, and VAT. Symbols and error bars indicate individual mice and mean values ± SD. Unpaired t-test: ns, not significant; *p ≤ 0.05, **p ≤ 0.01, ***p ≤ 0.001, ****p ≤ 0.0001. Data from two experiments, representative of at least three independent experiments (≥ 3 mice/group).

### Figure S9 (related to Figure 6F). Unstable VAT Treg cells in the 'Tmem' cluster of ΔpTreg mice

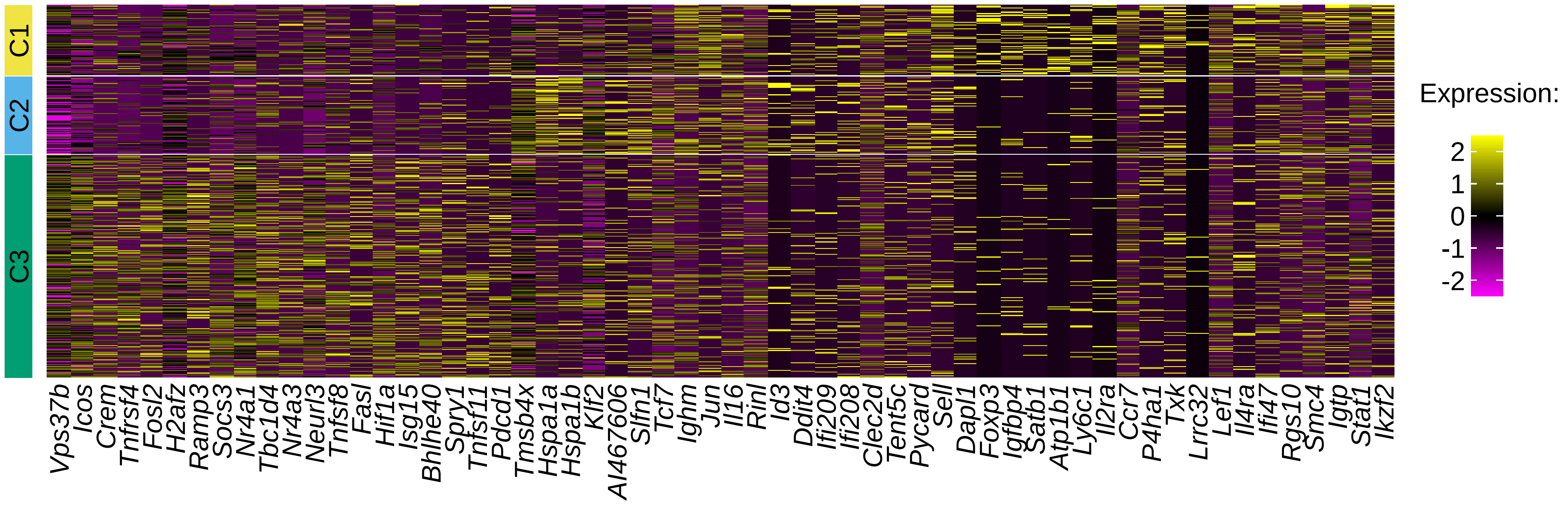


**Figure S9 (related to Figure 6F). Unstable VAT Treg cells in the 'Tmem' cluster of ΔpTreg mice**
Sub-clustering of the 'Tmem' cluster from scRNA/TCR-seq of VAT CD4$^{+}$ T cells in Foxp3$^{RFP/GFP}$ (WT), ΔtTreg, and ΔpTreg mice after 20 weeks of HFD (Figure 4D). Heatmap showing the top 20 differentially expressed genes (rows) per sub-cluster (top colors) in individual cells (columns) from Figure 6F. The top 50 differentially expressed genes are listed in Supplementary Table S2.

#### Figure S10 (related to Figure 7). Characterization of VAT ex-tTreg cells

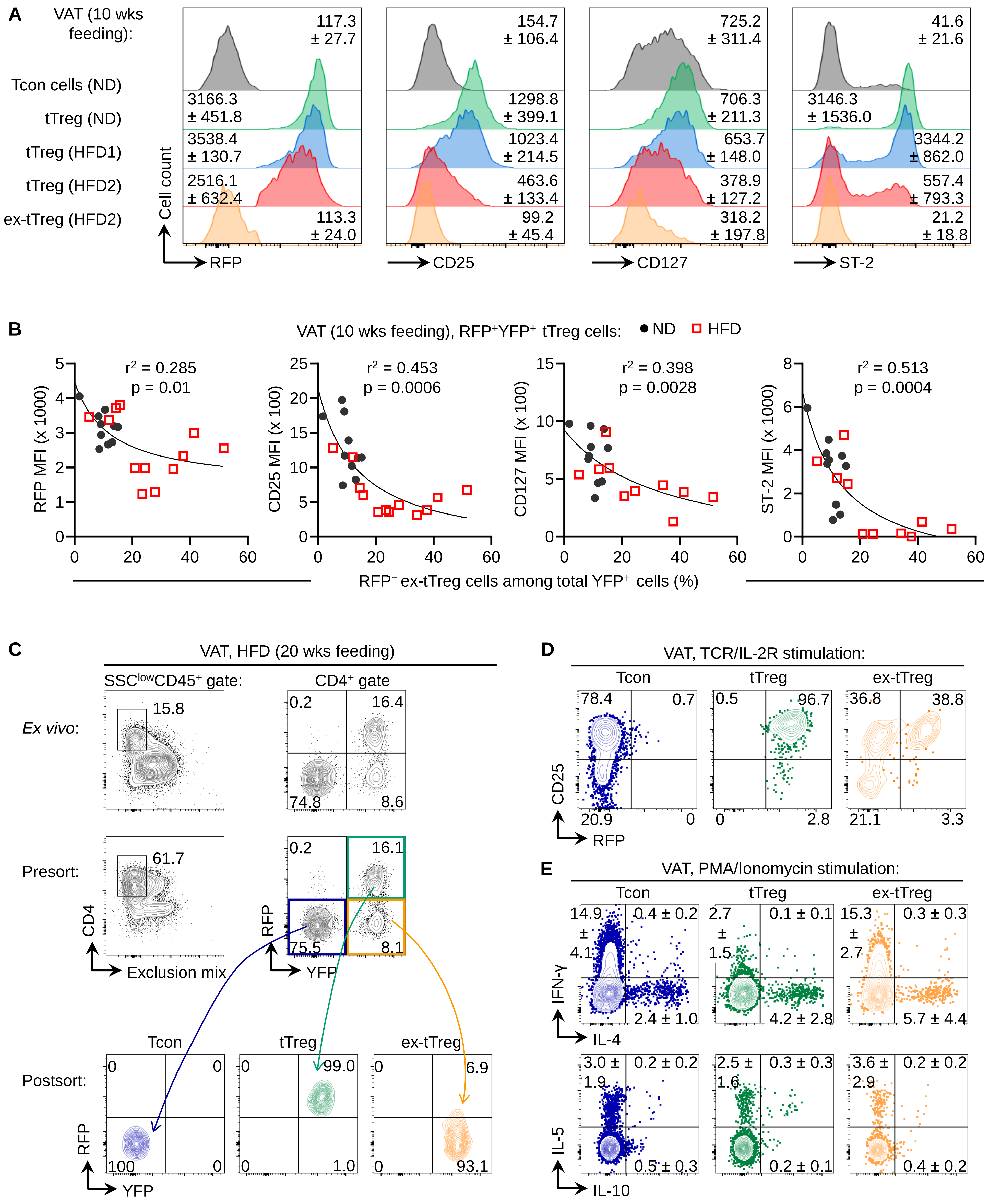

**Figure S10 (related to Figure 7). Characterization of VAT ex-tTreg cells**

(A and B) Flow cytometry of VAT CD4$^+$ T cell subsets in R26Y x Foxp3$^{RFP/GFP}$ mice fed ND or HFD for 10 weeks (Figure 7). (A) Histograms showing Foxp3-driven RFP (top, left) and selected surface markers of the VAT Treg cell signature in gated CD4$^+$ T cells. Mean MFI ± SD. (B) Correlation between marker expression in RFP$^+$YFP$^+$ tTreg cells from (A) and frequency of RFP$^-$ ex-tTreg cells among total YFP$^+$ cells in VAT. Symbols represent individual mice. Pearson correlation analysis ($r^2$ = coefficient of determination). Data from three independent experiments (n = 10 – 12/group).

(C-E) *In vitro* characterization of sorted VAT ex-tTreg cells from R26Y x Foxp3$^{RFP/GFP}$ mice fed HFD for 20 weeks. (C) Sorting strategy and post-sort analysis of ex-tTreg, Tcon, and tTreg cells. (D) Flow cytometry of CD25 vs. RFP expression following TCR/IL-2R stimulation (day 3) VAT of 6 mice were pooled. (E) Intracellular cytokine production following PMA/Ionomycin stimulation. Numbers indicate mean percentages ± SD. Data from a single experiment (n = 6).

**Figure S11 (related to Figure 4). Isolation of VAT CD4$^+$ T cells for scRNA/TCR-seq**

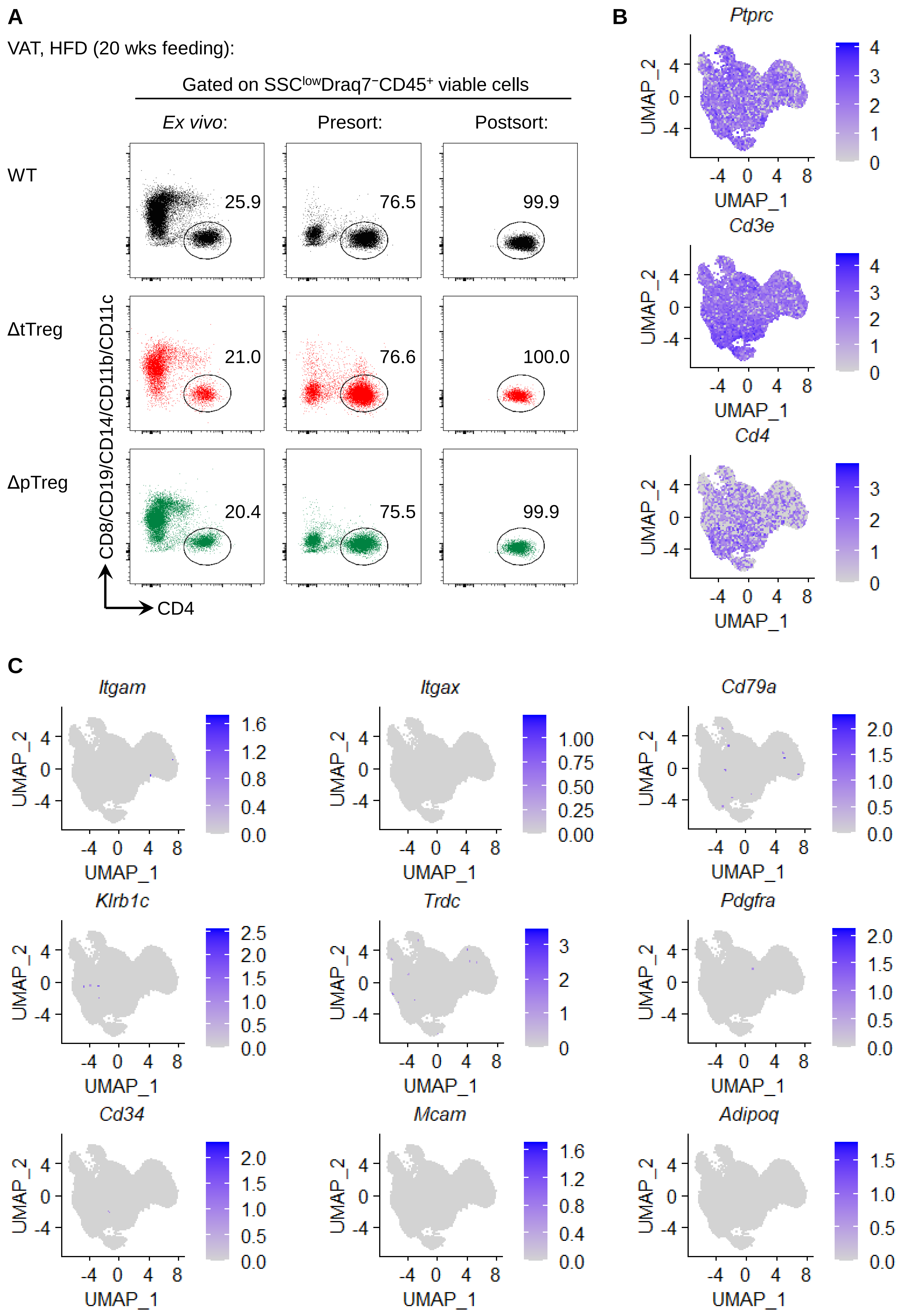

**Figure S11 (related to Figure 4). Isolation of VAT CD4$^+$ T cells for scRNA/TCR-seq**

(A) Sorting strategy. After 20 weeks of HFD feeding, pooled VAT single-cell suspensions from Foxp3$^{RFP/GFP}$ (WT, top; n = 5), ΔtTreg (middle; n = 4), and ΔpTreg (bottom; n = 4) mice were subjected to magnetic bead-based lineage depletion (CD8, CD19, CD14, CD11b, CD11c) and FACS purification (SSC$^{low}$Draq7$^-$CD45$^+$CD4$^+$). Purified cells were used for scRNA/TCR-seq.

(B and C) Validation of CD4$^+$ T cell identity in sequenced cells. UMAP projections show (B) *Ptprc* (CD45), *CD3e* (CD3ε), and *Cd4* (CD4) expression and (C) absence of lineage markers (*Itgam*, *Itgax*, *Cd79a*, *Klrb1c*, *Trdc*, *Pdgfra*, *Cd34*, *Mcam*, *Adipoq*) by scRNA-seq.

**Table S1.** Top 50 differentially expressed genes among VAT CD4$^+$ T cell clusters (see Figures 4D and S6 for details)

| | Treg | Tmem | Th2 | Th1/Tex | Th1 | Tex | Tn | CTL |
|---|---|---|---|---|---|---|---|---|
| 1 | Cxcl2 | S1pr1 | Hilpda | Ifng | Ly6c2 | Sostdc1 | Igfbp4 | Ccl5 |
| 2 | Areg | Il7r | Gadd45b | Pdcd1 | Nkg7 | Spp1 | Lef1 | Gzmk |
| 3 | Klrg1 | Rps27 | Nfkbia | Tigit | Epsti1 | Lag3 | Ccr7 | Nkg7 |
| 4 | Tnfrsf9 | Klf2 | Tnfsf11 | AW112010 | Tbx21 | Tnfsf8 | Dapl1 | Eomes |
| 5 | Ikzf2 | Slamf6 | Ramp3 | Lag3 | Ifngr1 | Cd200 | Ly6c1 | Ccr5 |
| 6 | Cd74 | Itgb1 | Atf3 | Tnfrsf4 | Crip1 | Gpm6b | Sell | Gm30211 |
| 7 | Foxp3 | Rflnb | Zfp36 | Il10 | Ctsw | Eea1 | Satb1 | Esm1 |
| 8 | Itgav | Gpm6b | Ltb4r1 | Cxcr6 | Ccl5 | Tbc1d4 | Atp1b1 | Crtam |
| 9 | Il1rl1 | Spry1 | Socs2 | Bhlhe40 | S100a4 | 1700019D03Rik | Actn1 | Nr4a2 |
| 10 | Il2ra | Tbc1d4 | Nfkbiz | Ctla4 | Fasl | Izumo1r | Rps20 | Igkc |
| 11 | Capg | Prkca | Ctla2a | Eea1 | Itgb1 | Slamf6 | Rps19 | Ctla2a |
| 12 | Ctla4 | Adk | Bcl2a1d | Maf | Lgals1 | Marcksl1 | Txk | Slamf7 |
| 13 | Pim1 | Lrig1 | Bcl2a1b | Ccr5 | Ahnak | Tox | Rps29 | Prf1 |
| 14 | Tnfrsf4 | Socs3 | Gata3 | Dusp1 | Zfp36l2 | Cd83 | Foxp1 | Ccl4 |
| 15 | Cd81 | Cd9 | Pim1 | Lilrb4a | Vim | Ptger2 | Rpl35a | Rpa2 |
| 16 | Ltb4r1 | Zc3hav1 | Il1rl1 | Ctsb | Dusp5 | Angptl2 | Rpl36a | Gpr183 |
| 17 | Tigit | Il6ra | Neurl3 | Aopep | Cxcr3 | Cacna1d | Npc2 | Ctsw |
| 18 | Sdc4 | Icos | Lmna | Tnfrsf18 | Ier2 | Ccdc28b | Pik3ip1 | Plek |
| 19 | Tnfrsf18 | Pou2f2 | Cdkn1a | Rgs16 | Hcst | Fam43a | Rps24 | Sh2d1a |
| 20 | Rln3 | Gramd3 | Ifrd1 | Coch | Il2rb | Ptpn11 | Rpl35 | Tspan3 |
| 21 | Gadd45b | Itga4 | Nfkbid | Sdf4 | Thy1 | Plagl1 | Pdlim4 | Cst7 |
| 22 | S100a10 | Sidt1 | Cd40lg | Asb2 | Ms4a4b | Tiam1 | Rps28 | Baiap3 |
| 23 | Fgl2 | Ehd3 | Ccrl2 | Serpina3g | Ubc | Prelid2 | Rplp0 | Fasl |
| 24 | Ccr8 | Atp1b3 | Ccl1 | Anxa2 | Ccnd3 | Il21 | Rflnb | Ap3s1 |
| 25 | Glrx | Gm8369 | Nr4a1 | Lilr4b | S100a6 | Cd82 | Rps7 | Ms4a4b |
| 26 | Dusp4 | Cd28 | Ccr2 | Lgals3 | Dennd4a | Ifi27l2a | Rps8 | St3gal6 |
| 27 | Cd83 | Gpr183 | Fosb | Ehd1 | Gramd3 | Mrps6 | Rpl12 | Cd27 |
| 28 | Samsn1 | Tspan13 | Tnfaip3 | Zap70 | Zyx | Nrn1 | Gpr18 | Sp100 |
| 29 | Nfkbia | Ripor2 | Samsn1 | S100a11 | Tagln2 | Pou2f2 | Rps4x | Myb |
| 30 | Pglyrp1 | Shld1 | Cish | Tbx21 | Dnajc15 | 2310001H17Rik | Rpl8 | Gpr18 |
| 31 | Zfp36l1 | Acot2 | Dusp5 | Arap2 | Btg2 | Hif1a | Rps26 | Ifitm10 |
| 32 | Traf1 | Cenpa | Jund | Podnl1 | Rora | Tnfaip8 | Pdk1 | Serpina3g |
| 33 | Il10ra | Rnf19a | Prr7 | Il21 | Cd226 | Creb3l2 | Rplp1 | Adam19 |
| 34 | Rel | Cdk11b | Tgfb1 | Sdcbp2 | Anxa2 | Raf1 | Rpl5 | Gtf2i |
| 35 | Dgat1 | Gm45552 | Fosl2 | Serpina3f | Rgs1 | Nfatc1 | Ifngr2 | Tox |
| 36 | Jund | Ssh2 | Ptpn13 | Nkg7 | Cxcr6 | Zap70 | Rps27a | Lax1 |
| 37 | Hopx | Samhd1 | Csrnp1 | Mmd | Lyst | Ctsb | Rpl10a | Ctsd |
| 38 | Nfkbiz | Nrp1 | Ell2 | Dusp14 | Anxa6 | Cd9 | Rps16 | Sh2d2a |
| 39 | H2afz | Gm2682 | Rab4a | Ctsd | Icos | S100a11 | Limd2 | Zyx |
| 40 | Gata3 | Crlf3 | Cxcr6 | Tnfsf8 | Ifng | Rilpl2 | Rpl28 | Runx3 |
| 41 | Izumo1r | Cdkn1b | Ier5 | S100a6 | Adgre5 | Smco4 | Dgka | Plcxd2 |
| 42 | Nfkbid | D8Ertd738e | Ern1 | Furin | Serpinb9 | Ramp1 | Rpl13 | Sart3 |
| 43 | Zfp36 | Dusp10 | Gm42031 | Glrx | Hmgb2 | Tpi1 | Rps27 | Txk |
| 44 | Anxa1 | Slc4a7 | Ldha | Batf | Abcb1b | Ddit4 | A930005H10Rik | Prkch |
| 45 | Nr4a1 | Stk38 | Hdc | Spp1 | Rcbtb2 | Cst7 | Nsg2 | Fyn |
| 46 | Ctla2a | P2ry10 | Areg | Rgs1 | Dusp2 | Stx11 | Cmah | Armc7 |
| 47 | Atf3 | Rasa3 | Rgs2 | Serpinb1a | Socs2 | Smpdl3a | Evl | Tapbpl |
| 48 | Hilpda | Sap18 | Calca | Ccr8 | Egr1 | Prkca | Selenop | Ier3 |
| 49 | Penk | Arhgap31 | Rgcc | Trav13d-1 | Tnf | Cd247 | Gm26917 | Rgs16 |
| 50 | Lmna | Cblb | Rrad | Ccl4 | Cd69 | Penk | Tubb2a | Trbv3 |

**Table S2.** Top 50 differentially expressed genes among 'Tmem' sub-clusters (see Figures 6F and S9 for details)

| C1 | | | | C2 | | | | C3 | | | |
|---|---|---|---|---|---|---|---|---|---|---|---|
| 1 | Foxp3 | 26 | Gbp2 | 1 | Hspa1a | 26 | Evl | 1 | Ramp3 | 26 | Cdk11b |
| 2 | Igfbp4 | 27 | Tgtp1 | 2 | Hspa1b | 27 | Tubb5 | 2 | Tnfrsf4 | 27 | Gpm6b |
| 3 | Ly6c1 | 28 | Gadd45b | 3 | AI467606 | 28 | Tbc1d10c | 3 | Crem | 28 | Gna13 |
| 4 | Ccr7 | 29 | Ccl5 | 4 | Klf2 | 29 | 4930453N24Rik | 4 | Fosl2 | 29 | Bcl2a1d |
| 5 | Il2ra | 30 | Dennd4a | 5 | Jun | 30 | Gm26917 | 5 | Icos | 30 | Ubc |
| 6 | Satb1 | 31 | Trbv31 | 6 | Slfn1 | 31 | Arhgap45 | 6 | Pdcd1 | 31 | Junb |
| 7 | Igtp | 32 | Ecm1 | 7 | Ighm | 32 | Zfp36 | 7 | Tnfsf8 | 32 | Bcl2a1b |
| 8 | Sell | 33 | Ift80 | 8 | Tcf7 | 33 | Arrb2 | 8 | Bhlhe40 | 33 | Rora |
| 9 | Stat1 | 34 | Gbp4 | 9 | Il16 | 34 | 1810009A15Rik | 9 | Fasl | 34 | Emb |
| 10 | P4ha1 | 35 | Gpr18 | 10 | Id3 | 35 | Fam189b | 10 | Nr4a3 | 35 | Dusp10 |
| 11 | Lef1 | 36 | Selplg | 11 | Dapl1 | 36 | Gmfg | 11 | Vps37b | 36 | Gpr132 |
| 12 | Ikzf2 | 37 | Cyth4 | 12 | Sell | 37 | Ppp1r18 | 12 | Tbc1d4 | 37 | Gramd3 |
| 13 | Smc4 | 38 | Irf1 | 13 | Ifi209 | 38 | S100a10 | 13 | Tnfsf11 | 38 | Cstb |
| 14 | Atp1b1 | 39 | Tubb2a | 14 | Ddit4 | 39 | Slc25a3 | 14 | Socs3 | 39 | Prkca |
| 15 | Jun | 40 | Npc2 | 15 | Pycard | 40 | Clint1 | 15 | H2afz | 40 | Cwc25 |
| 16 | Txk | 41 | Ighm | 16 | Tent5c | 41 | Zc3h12a | 16 | Hif1a | 41 | Sap18 |
| 17 | Ifi47 | 42 | Resf1 | 17 | Rinl | 42 | Brd2 | 17 | Isg15 | 42 | Dusp5 |
| 18 | Lrrc32 | 43 | Sesn1 | 18 | Tmsb4x | 43 | Capg | 18 | Spry1 | 43 | Eea1 |
| 19 | Il4ra | 44 | Actn1 | 19 | Ifi208 | 44 | Bysl | 19 | Neurl3 | 44 | Arpc3 |
| 20 | Rgs10 | 45 | Ptp4a3 | 20 | Clec2d | 45 | Mat2a | 20 | Nr4a1 | 45 | Odc1 |
| 21 | Arl5c | 46 | Plcxd2 | 21 | Ptprcap | 46 | Plscr1 | 21 | Gch1 | 46 | Ptpn11 |
| 22 | Trat1 | 47 | Pde4b | 22 | Ltb | 47 | Lmna | 22 | Got1 | 47 | D8Ertd738e |
| 23 | Pim1 | 48 | Aopep | 23 | St6gal1 | 48 | Chd1 | 23 | Tigit | 48 | Areg |
| 24 | Cd74 | 49 | Gm14085 | 24 | Fam78a | 49 | Selenok | 24 | P2ry10 | 49 | Zc3hav1 |
| 25 | Ifngr2 | 50 | Itm2a | 25 | Dennd2d | 50 | Suco | 25 | Rnf125 | 50 | Itgb1 |
